# The Pif1 helicase is actively inhibited during meiotic recombination which restrains gene conversion tract length

**DOI:** 10.1101/2021.01.21.427607

**Authors:** Dipti Vinayak Vernekar, Giordano Reginato, Céline Adam, Lepakshi Ranjha, Florent Dingli, Marie-Claude Marsolier, Damarys Loew, Raphaël Guérois, Bertrand Llorente, Petr Cejka, Valérie Borde

## Abstract

Meiotic recombination ensures proper chromosome segregation to form viable gametes and results in gene conversions events between homologs. Conversion tracts are shorter in meiosis than in mitotically dividing cells. This results at least in part from the binding of a complex, containing the Mer3 helicase and the MutLβ heterodimer, to meiotic recombination intermediates. The molecular actors inhibited by this complex are elusive. The Pif1 DNA helicase is known to stimulate DNA polymerase delta (Pol δ) -mediated DNA synthesis from D-loops, allowing long synthesis required for break-induced replication. We show that Pif1 is also recruited genome wide to meiotic DNA double-strand break (DSB) sites. We further show that Pif1, through its interaction with PCNA, is required for the long gene conversions observed in the absence of MutLβ recruitment to recombination sites. *In vivo*, Mer3 interacts with the PCNA clamp loader RFC, and *in vitro*, Mer3-MutLβ ensemble inhibits Pif1-stimulated D-loop extension by Pol δ and RFC-PCNA. Mechanistically, our results suggest that Mer3-MutLβ may compete with Pif1 for binding to RFC-PCNA. Taken together, our data show that Pif1’s activity that promotes meiotic DNA repair synthesis is restrained by the Mer3-MutLβ ensemble which in turn prevents long gene conversion tracts and possibly associated mutagenesis.

## Introduction

Meiosis is a specialized cell division used by sexually reproducing organisms to produce haploid gametes from a diploid parent cell. Homologous recombination during the first meiotic division prophase forms crossovers (COs) that physically connect the homologs, which ensures their proper segregation during the reductional division. Defects in these processes result in aneuploidy and infertility. Meiotic recombination begins with genome-wide programmed DNA double-strand breaks (DSBs) catalyzed by Spo11. These DSBs are further processed by end resection, producing 3’ ends that preferably invade the homologous chromosome as a repair template, resulting in the formation of a D-loop intermediate. D-loops are further processed to either COs or non-crossover (NCOs) products. COs are majorly formed through the ZMM (Zip-Msh-Mer) pathway, which includes eight conserved proteins, Zip1-4, Spo16, Mer3, Msh4 and Msh5 that are proposed to stabilize the D-loop intermediates (1,2).

Both COs and NCOs involve formation of heteroduplex DNA during DSB repair that may result in gene conversions after mismatch repair. Gene conversions influence genetic diversity by promoting allelic transmission distortion, hence it is necessary to control their extent, but the regulatory factors are not fully known (3). In the budding yeast *Saccharomyces cerevisiae*, one of the three MutL heterodimers, MutLβ (Mlh1-Mlh2), along with Mer3, a meiosis specific helicase and a ZMM member, is known to limit the extent of meiotic gene conversions (4). MutLβ is recruited to DSB hotspots through its interaction with Mer3. The meiotic functions of MutLβ depend on its interaction with Mer3, which is lost in a *mer3R893E* point mutant. MutLβ also specifically binds D-loop structures *in vitro* but lacks endonuclease activity, contrary to the other MutL heterodimers (4,5). The analysis of heteroduplex DNA tracts lengths in a SK1/S288C hybrid background revealed that a *mlh2Δ* and *mer3R893E* single mutants have longer tracts than the reference at both CO and NCO sites. In this hybrid diploid, the *mlh2Δ* mutant still preferentially uses the ZMM pathway for CO formation like wild-type cells but shows reduced spore viability (4). As MutLβ does not have any enzymatic activity, it was proposed that the Mer3-MutLβ complex acts as a physical barrier to the proteins involved in extending D-loops during meiotic DNA repair synthesis. The proteins involved may comprise the DNA polymerase(s) themselves, but also other members of the replication machinery or helicases (6,7).

Pif1 is a highly conserved 5’ to 3’-directed DNA helicase having multiple functions in mitochondria and in the nucleus. Pif1 has two isoforms, mitochondrial and nuclear, which are generated by using the first and the second start codons in *PIF1* mRNA, respectively, and can be separately studied thanks to mutations that affect the use of these start codons (8). Pif1 was found to inhibit telomere lengthening and *de novo* telomere addition (8), which was further confirmed by the discovery that it is a negative regulator of telomerase *in vitro* (9). In addition, major functions of Pif1 include mitochondrial genome maintenance, processing of Okazaki fragments, resolution of G-quadruplex (G4) structures and DNA synthesis during break-induced replication (BIR) (10).

In BIR, cells employ Pol δ to synthesize DNA over very long genomic distances, by strand-displacement synthesis and bubble migration (11–13). The continuous DNA length synthesized by Pol δ in BIR is much longer than during S phase, where it catalyses the synthesis of Okazaki fragments on the lagging strand (13,14). Contrary to the leading strand DNA polymerase epsilon, Pol δ does not interact with the MCM helicase and is therefore thought to be less processive (15). During BIR, Pif1 is enriched at the DSB site and along the repair template molecule, and complementarily, cells lacking nuclear Pif1 are deficient in BIR and show a decreased recruitment of Pol δ (12,13). Besides BIR, experiments in vegetatively growing cells have shown that Pif1 is also important for both long conversion tracts and for crossovers during repair of a site-specific HO DSB by homologous recombination (13). In agreement with these *in vivo* results, *in vitro*, Pol δ-mediated DNA synthesis is stimulated and produces longer extension products in the presence of Pif1 (13). DNA unwinding thus allows extending DNA synthesis in the migrating D-loop structures (13). Indeed, in the absence of Pif1, recombination intermediates are not extended further than 1kb (13). *In vitro*, Pif1 does not physically interact with Pol δ, but interacts through its C-terminal region with the essential cofactor polymerase sliding clamp, PCNA (Proliferating Cell Nuclear Antigen) (16). A quadruple point mutant in the C terminus of Pif1 (*pif1R3E*) looses the interaction with PCNA, produces shorter extension products from D-loops as compared to wild type Pif1 *in vitro*, and *in vivo*, decreases BIR efficiency (16).

The properties of Pif1 to facilitate DNA synthesis in D-loop structures prompted us to check its involvement in the longer gene conversion events seen in *mlh2Δ* cells during meiosis. We found that Pif1 associates with DSB hotspots during meiosis. Longer gene conversion events observed in *mlh2Δ* cells require functional nuclear Pif1 and are dependent on the Pif1-PCNA interaction. Furthermore, Mer3 interacts *in vivo* with Rfc1 and Pif1, and *in vitro*, the Mer3-MutLβ complex inhibits Pif1 function to promote Pol δ mediated D-loop extension. Our results suggest that Mer3-MutLβ complex inhibits Pif1 activity during meiotic DNA repair synthesis resulting in limiting the lengths of gene conversions.

## Materials and Methods

### Yeast manipulations

All yeast strains are derivatives of the SK1 background unless otherwise stated (genotypes in Table S1). All experiments were performed at 30°C. For synchronous meiosis induction, cells containing the *pCUP1-IME1* construct were used and meiosis induced as described (17,18). For strain constructions and spore viability measurements, sporulation was performed on solid sporulation medium for two days. For octad sequencing, sporulation of S288CxSK1 hybrids was performed in liquid cultures as described (4).

### Construction of yeast strains

Yeast strains were obtained by direct transformation or crossing to obtain the desired genotype. Site-directed mutagenesis, C-terminal tag insertions and gene deletions were introduced by PCR. All transformants were validated using PCR to discriminate between correct and incorrect integrations, and sequenced to ensure epitope tag insertion or mutagenesis. Mer3 and Pif1 were C-terminally tagged with the TAP tag sequence (19). Rfc1-TAP was described previously (20). For Mer3, a GGGGSGGGGS linker was added between Mer3 and the TAP sequence, as done previously for Mer3-Flag, to preserve spore viability (4). Pif1 was tagged with 13 copies of the Myc epitope (21). *pif1m2* was introduced by a 2-step strategy using pJL73 (22). The *pif1R3E* (I817R M820R L821R R823E) mutation was introduced by CRISPR-Cas9 mediated cleavage, as described (4).

### Chromatin immunoprecipitation, real-time quantitative PCR and ChIP-seq

For ChIP-qPCR or ChIP-seq analysis of Pif1-Myc, 2.5 × 10^8^ or 1.2 × 10^9^ cells, respectively, from p*CUP1-IME1* synchronized time-courses were processed at t = 0 h (time of meiosis induction) or at t = 5 h exactly as described (18,23).

Quantitative PCR was performed from the immunoprecipitated DNA or the whole cell extract using a 7900HT Fast Real-Time PCR System and SYBR Green PCR master mix (Applied Biosystems, Thermo Scientific) as described (23). Results were expressed as % of DNA in the total input present in the immunoprecipitated sample and normalized first by a negative control site on chromosome III, *LDLR* (24) and then by the 0 h time-point. Primers for *GAT1*, *BUD23*, *HIS4LEU2*, Axis have been described (4).

### Illumina sequencing of ChIP DNA and read normalization

For ChIP-seq, immunoprecipitated DNA was purified and analyzed by sequencing as described (18) on a Illumina HiSeq2500 apparatus, generating paired-end 50 bp reads. Pif1-Myc13 ChIP-seq was performed in duplicate at t = 5 h in meiotic cells (VBD1907) synchronized with the copper-inducible system. To obtain the DSB-specific Pif1 ChIP-seq signal, the negative control was Pif1-Myc13 ChIP-seq in a DSB-deficient *spo11* strain (VBD1930) at the same time-point, to reveal only DSB-specific Pif1 binding sites. For the analysis presented in Figure S1A, the Pif1-Myc13 signal from VBD1907 (*SPO11*) or VBD1930 (*spo11*) was subtracted by the Mlh3-Myc ChIP-seq signal from a *spo11* strain, in which Mlh3 does not associate with chromatin (25). Bioinformatic analyses for alignment, reads normalization, smoothing were done as described before, using custom R and Python scripts as well as the Galaxy (www.galaxy.org) platform (18,26).

### Genome-wide meiotic recombination events inferred from octad analysis

Genomic DNA from the eight meiotic products of single meioses was prepared using Qiagen Genomic-tip 100 kit, and sequenced using an Illumina HiSeq 2500 instruments. We studied two octads of each genotype (*pif1m2*, *pif1m2 mlh2Δ* and *pif1R3E mlh2Δ*). Sequencing reads from octad were aligned on the S288C and SK1 reference sequences to genotype SNPs and deduce recombination events exactly as described (4).

### Label-free mass spectrometry analysis of Mer3-TAP complexes

1 liter (2.5×10^10^) of cells at 5 h of a p*CUP1-IME1* synchronized meiosis from Mer3-TAP strain (VBD1875) or untagged Mer3 strain (VBD2119) as a negative control were used. Each condition was made in four independent replicates. PMSF (phenylmethylsulfonyl fluoride) to a final concentration of 1 mM was added to the culture prior harvesting cells. Cells were washed with TAP lysis buffer (50 mM Tris/HCl pH 7.5; 1 mM EDTA; 0.5% NP-40; 10% glycerol; 300 mM NaCl; 1 mM PMSF; 1X Complete Mini EDTA-Free (Roche); 1X PhosSTOP (Roche)), resuspended in about 2 ml of the same buffer and frozen as noodles in liquid nitrogen. For lysis, cells were ground with a 6775 Freezer/Mill cryogenic grinder (SPEX SamplePrep). The resulting powder was resuspended in 50 ml TAP Lysis buffer plus 1 mM PMSF and 1X Complete EDTA-free protease inhibitor cocktail (Roche). The lysate was cleared by centrifugation at 4000 g for 10 min and then incubated with 1 ml (packed volume 600 μl) of lgG Sepharose beads (GE Healthcare) for 1 h at 4 °C. The beads were washed three times with TAP lysis buffer and resuspended in the same buffer to 1 ml total. 60 μl packed beads were directly processed for mass spectrometry analysis.

Proteins on beads were washed twice with 100 μl of 25 mM NH4HCO3 and subjected to on-beads digestion with 0.2 μg of trypsine/LysC (Promega) for 60 min in 100 μl of 25 mM NH4HCO3. Sample were then loaded onto a homemade C18 StageTips for desalting. Peptides were eluted using 40/60 MeCN/H2O + 0.1% formic acid and vacuum concentrated to dryness. Prior analyses, digests were reconstituted in 10 μL of 0.3% TFA in 2/98 MeCN/H2O and 5 μL were analyzed by LC-MS/MS using an RSLCnano system (Ultimate 3000, Thermo Scientific) interfaced on-line to an Orbitrap Fusion Tribrid mass spectrometer (Thermo Scientific) as in (27). For identification, data were searched against the *S. cerevisiae* UniProt database (UP000002311, taxonomyID 559292) using SEQUEST-HT through proteome discoverer (version 2.2). Enzyme specificity was set to trypsin and a maximum of two-missed cleavage sites was allowed. Oxidized methionine and N-terminal acetylation were set as variable modifications. Maximum allowed mass deviation was set to 10 ppm for monoisotopic precursor ions and 0.6 Da for MS/MS peaks. The resulting files were further processed using myProMS v3.9 (28). For identification, FDR calculation used Percolator (Spivak et al., 2009) and was set to 1% at the peptide level for the whole study. The label free quantification was performed by peptide Extracted Ion Chromatograms (XICs) computed with MassChroQ v2.2.1 (29). For protein quantification, XICs from proteotypic peptides shared between compared conditions (TopN matching) with missed cleavages were used. Median and scale normalization was applied on the total signal to correct the XICs for each biological replicate (n=4). To estimate the significance of the change in protein abundance, a linear model (adjusted on peptides and biological replicates) based on two-tailed T-test was performed and p-values were adjusted with a Benjamini–Hochberg FDR procedure. Proteins with at least two-fold enrichment, adjusted p-value < 0.05, and at least three distinct peptides in all replicates were considered significantly enriched in sample comparisons.

### Coimmunoprecipitation and Western blot analysis

1.2×10^9^ cells were harvested, washed once with PBS, and lyzed in 3 ml lysis buffer (20 mM HEPES/KOH pH7.5; 150 mM NaCl; 0.5% Triton X-100; 10% Glycerol; 1 mM MgCl2; 2 mM EDTA; 1 mM PMSF; 1X Complete Mini EDTA-Free (Roche); 1X PhosSTOP (Roche); 125 U/mL benzonase) with glass beads three times for 30 s in a Fastprep instrument (MP Biomedicals, Santa Ana, CA). The lysate was incubated 1 h at 4°C. 100 μL of PanMouse IgG magnetic beads (Thermo Scientific) were washed with 100 μL lysis buffer, preincubated in 100 μg/ml BSA in lysis buffer for 2 h at 4°C and then washed twice with 100 μL lysis buffer. The lysate was cleared by centrifugation at 13,000 × g for 5 min and incubated overnight at 4°C with washed PanMouse IgG magnetic beads. The magnetic beads were washed four times with 1 mL of wash buffer (20 mM HEPES/KOH pH7.5; 150 mM NaCl; 0.5% Triton X-100; 5% Glycerol; 1 mM MgCl2; 2 mM EDTA; 1 mM PMSF; 1X Complete Mini EDTA-Free (Roche); 1X Phos-STOP (Roche)). The beads were resuspended in 30 μL of TEV-C buffer (20 mM Tris/HCl pH 8; 0.5 mM EDTA; 150 mM NaCl; 0.1% NP-40; 5% glycerol; 1 mM MgCl2; 1 mM DTT) with 4 μl TEV protease (1 mg/ml) and incubated for 2 h at 23°C under agitation. The eluate was transferred to a new tube. Beads eluate was heated at 95°C for 10 min and loaded on polyacrylamide gel (4-12% Bis-Tris gel (Invitrogen)) and run in MOPS SDS Running Buffer (Life Technologies). Proteins were then transferred to PVDF membrane using XCell II^TM^ Blot Module (ThermoFisher Scientific) at 25 V constant for 2 h. Proteins were detected using c-Myc mouse monoclonal antibody (9E10, Santa Cruz, 1/500), Flag mouse monoclonal antibody (M2, Sigma, 1/1000) or TAP rabbit monoclonal antibody (CAB1001, Invitrogen, 1/2000). The TAP antibody still detects the CBP (Calmodulin Binding Protein) moiety after TEV cleavage of the TAP tag. Signal was detected using the SuperSignal West Pico or Femto Chemiluminescent Substrate (ThermoFisher). Signal was quantified after image acquisition with Chemidoc system (Biorad).

### Protein purification

The purification procedure for Mer3 helicase-dead (Mer3 K167A or Mer3-hd) and MutLβ was as described (4). Yeast RPA was expressed in BL21 (DE3) pLysS cells (Promega) from p11d-scRPA vector (a kind gift from M. Wold, University of Iowa) and purified as described for human recombinant RPA (30). PCNA and RFC were expressed in *E. coli* and purified according to previously established procedures (31,32). The three-subunit Pol δ was expressed in the yeast train WDH668 as described previously (33) and purified according to existing protocols (13,34). The sequences for the expression of FLAG-Pif1 (aa 40-859) and FLAG-Rad54 were codon-optimized for the expression in *E. coli* and obtained from GenScript. The genes were cloned into pMALT-P (Kowalczykowski laboratory) using *Nde*I and *Pst*I restriction sites. The proteins were expressed in BL21 (DE3) pLysS cells (Promega) induced with 0.5 mM isopropyl-1-thio β-D-galactopyranoside (IPTG) for 16 h at 18°C (2l for each construct). After collection, cells were resuspended and sonicated in 300 mM NaCl Buffer A (50 mM Tris-HCl pH 7.5, 1 mM PMSF, 0.5 mM βME, 10% Glycerol and 1 mM EDTA) supplemented with 1:500 Protease Inhibitor cocktail (Sigma, P8340), 30 μg/ml Leupeptin (Merk Millipore) and 0.1% NP40. The lysate was clarified by centrifugation and the soluble extract was incubated with M2 anti-FLAG affinity resin (Sigma) at 4°C for 1 h. The resin was washed with 300 mM Buffer A supplemented with 0.1% NP40. The salt concentration was decreased with a wash step to 150 mM NaCl first with and then without 0.1% NP40. The protein was eluted using Buffer A with 150 mM NaCl and 200 μg/ml 3XFLAG peptide (Sigma). Peak fractions were pooled, aliquoted, snap-frozen in liquid nitrogen and stored at −80 °C. The yield was ∼40 μg for Pif1 and ∼25 μg for Rad54 from 2 l of culture each. For Rad51, the codon-optimized sequence for bacterial expression was ordered from GenScript and cloned into pMALT-P using *Bam*HI and *Pst*I. The resulting vector expresses the Rad51 protein with a N-terminal MBP-tag cleavable by PreScission Protease. The protein was expressed as described above. The cell pellet from 2 l was resuspended and sonicated in 500 mM NaCl Buffer A (50 mM Tris-HCl pH 7.5, 1 mM PMSF, 1 mM DTT, 10% Glycerol) supplemented with 1:500 Protease Inhibitor cocktail (Sigma, P8465). The soluble extract obtained by centrifugation was supplemented with 0.01% NP40 and incubated with amylose resin (New England Biolabs) at 4°C for 1 h. The resin was washed with Buffer A with 1 M NaCl and subsequently with Buffer A with 300 mM NaCl but without PMSF. The protein was eluted in 300 mM NaCl Buffer A without PMSF supplemented with 10 mM maltose. Peak fractions were pooled and incubated with PreScission Protease (1:7, w/w) for 1 h at 4°C. The cleaved protein was diluted with 20 mM Tris-HCl pH 7.5 to reach 150 mM NaCl and loaded onto pre-equilibrated HiTrap Q HP column (GE Healthcare). The column was washed with 200 mM NaCl Buffer R (20 mM Tris-HCl pH 7.5, 1 mM EDTA, 0.5 mM DTT and 10% Glycerol). The protein was eluted with a salt gradient up to 700 mM NaCl in the same buffer. Peak fractions were pooled and dialyzed overnight against 100 mM NaCl Buffer R without EDTA. The dialyzed sample was aliquoted, snap-frozen in liquid nitrogen and stored at −80°C. The yield was ∼2 mg from 2 l of culture.

### Biochemical assays

The procedure to prepare the D-loop substrate for the helicase assay and the DNA binding experiments was described before (35) with minor modifications.

Helicase assays were performed in 15 μl of reaction buffer containing 35 mM Tris-HCl pH 7.5, 1 mM DTT, 7 mM MgCl, 0.1 mg/ml BSA, 2 mM phosphoenolpyruvate (PEP), 2 mM ATP and 80 U/ml Pyruvate kinase (Sigma). 1 nM of the D-loop substrate (in molecules) was incubated with 65.55 nM RPA and the indicated proteins for 30 min at 30°C. The reactions were then stopped with 5 μl of stop solution (150 mM EDTA, 2% SDS, 30% glycerol, 0.25% bromophenol blue) and 1 μl Proteinase K (14–22 mg/ml, Roche). Stopped reactions were incubated for 10 min at 30°C. The products were separated on native 10% polyacrylamide gels (acrylamide: bisacrylamide 19:1, Biorad). The gels were then dried and exposed to storage phosphor screens (GE Healthcare). The exposed screens were scanned using Typhoon FLA 9500 (GE Healthcare) and quantitated by ImageJ software.

DNA binding assays were carried out in binding buffer containing 1 nM substrate (in molecules), 35 mM Tris-HCl pH 7.5, 1 mM DTT, 7 mM MgCl, 0.1 mg/ml BSA and 2 mM ATP. The indicated proteins were added on ice and the reactions were incubated for 30 min at 30°C. After incubation, the reactions were supplemented with 5 μl of loading dye (50% glycerol and 0.25% bromophenol blue) and separated on native 4% polyacrylamide gels (acrylamide: bisacrylamide 19:1). Gels were processed and quantitated as described above.

D-loop extension assays were performed as described previously (13) with minor modifications. Briefly, the 90-mer oligonucleotide substrate (2.4 μM in nucleotides) was incubated with Rad51 (533 nM) in reaction buffer (final concentrations: 35 mM Tris-HCl pH 7.5, 1 mM DTT, 7 mM MgCl, 0.1 mg/ml BSA, 2 mM PEP, 2 mM ATP, dNTPs 100 μM each and 80 U/ml Pyruvate kinase) for 10 min at 37°C. To visualize the extended D-loop, 80 nCi/μl (final concentration) of [α-^32^P]-dCTP (Perkin Elmer) was added to the reaction. The reaction was then supplemented with RPA (400 nM) and further incubated for 5 min at 37°C. After the incubation Rad54 (40 nM) was added and incubation continued for 2 min at 23°C. pUC19 plasmid donor (36 μM in base pairs) was incorporated and incubation proceeded for 2 min at 30°C. RFC and PCNA (50 nM each) were added and the reaction was placed on ice for 2 min. Pol δ (40 nM), Pif1 (20 nM) and the indicated concentration of Mer3-hd and MutLβ were added and the reaction was incubated for 15 min at 15°C. Reactions were stopped with 5 μl of stop solution (150 mM EDTA, 2% SDS, 30% glycerol, 0.25% bromophenol blue) and 1 μl Proteinase K, and incubated for 10 min at 37°C. The products were separated by 1% agarose gel electrophoresis. Gels were dried and processed as described above. DNA extension reactions (15 μl) were carried out as described previously (34), using an oligonucleotide with 30 nt flap annealed to a plasmid based circular ssDNA (3197 nt) as a substrate. The reaction contained 25 mM Tris-acetate pH 8.5, 10 mM magnesium acetate, 125 mM NaCl, 1 mM ATP, 1 mM DTT, 0.1 mg/ml BSA, 1 mM PEP, 80 U/ml pyruvate kinase, 100 μM dNTPs (each), 100 ng DNA substrate, 20 nM PCNA, 20 nM RFC, 1 μM RPA (concentration saturating 100% ssDNA). The reaction was assembled on ice and pre-incubated for 1 min at 30^°^C. Next, 5 nM Pol δ and indicated amounts of Mer3-hd and MutLβ respectively were added and the reaction was incubated at 30^°^C for given time points before termination by the addition of 5 μl of STOP buffer (30 mM EDTA, 2% SDS, 30% glycerol, and 1 mg/ml bromophenol blue) and 1 μl proteinase K (14–22 mg/ml, Roche). The reaction was further incubated for 15 mins at 30^°^C for deproteination. The extension products were analysed on 1% agarose gel containing GelRed (1:10,000 v/v, Biotium) in TAE buffer. The gels were imaged by gel imager (InGenius3, GeneSys), the bands were quantitated using ImageQuant (GE Healthcare) and expressed as percentage of total DNA synthesis (extension) for each protein combination.

## Results

### Pif1 occupies meiotic DSB hotspots genome wide

To find proteins involved in producing longer gene conversions when the loading of MutLβ is lost, we tested the involvement of the Pif1 helicase. If Pif1 promotes longer gene conversions in the absence of MutLβ loading, Pif1 should be present at DSB hotspots. Hence, we first tested whether Pif1 was recruited to DSB hotspots during meiosis. We performed chromatin immunoprecipitation followed by qPCR analysis from synchronous meiotic cultures of Myc-tagged Pif1 cells at 5 h in meiosis, the expected time of meiotic DSB repair (36). Pif1-Myc was associated with the analyzed DSB hotspots in wild-type and *mlh2Δ* cells but not in the *spo11Y135F* mutant, indicating a DSB-dependent recruitment of Pif1 to recombination sites (Figure 1A). The deletion of *MLH2* did not modify enrichment of Pif1 to DSB sites, suggesting that Pif1 recruitment to DSB hotspots occurs independently of MutLβ.

**Figure 1:**
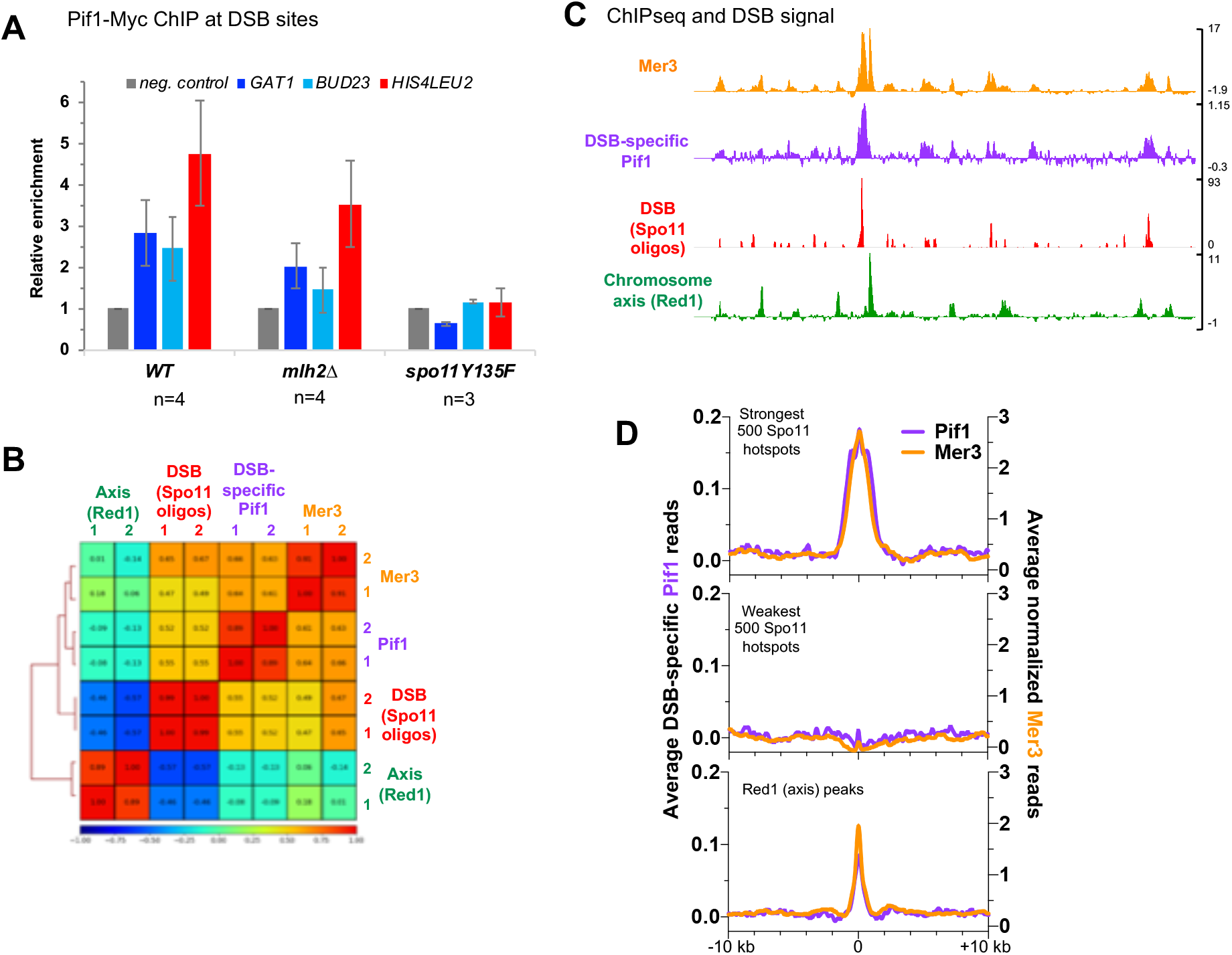
Pif1 occupies meiotic DSB hotspots genome wide. (A) Pif1-Myc levels at the three indicated meiotic DSB hotspots relative to a negative control site (LDLR) assessed by ChIP and qPCR at the 5 h time-point of meiotic time-courses. The signal is further normalized to the signal observed at t = 0 h. Values are the mean ± S.E.M. from the indicated number of independent experiments. (B) Correlation heatmap between DSBs, Red1, Mer3 and Pif1 ChIP-seq signals. For each replicate, normalized binding data of the indicated protein were used after smoothing with a 2000 bp window. Mer3 and Spo11 oligo data are from (25) and Red1 data are from (39). The comparison was made on the regions encompassing the Red1 peaks (933 peaks) (39), the strongest 1000 Spo11 oligo hotspots (44) and the strongest 1000 Mlh3 peaks (25). The Spearman correlation coefficient is indicated for each pair-wise comparison. (C) ChIP-seq analysis of Pif1 binding compared to the binding of the Mer3 (25), to DSBs (Spo11 oligos (25)), and to chromosome axis attachment sites (Red1 ChIP-seq (39)). Normalized data are smoothed with a 200 bp window. (D) Average ChIP-seq signal at the indicated features. Same data as in (C). The Mer3 signal is aligned on the indicated Spo11 hotspots midpoints from (44), and the Pif1 ChIP-seq signal on the p*CUP1-IME1* Spo11 hotspots midpoints (25), or on the summit of the Red1 peaks summits (39).

Pif1 was previously reported to associate with difficult to replicate regions such as G4-prone sequences (37) and tRNA genes (38). To further explore Pif1 distribution during meiosis, we performed ChIP-seq analysis for Pif1-Myc from cells at 5 h in meiosis. Independently of meiotic DSB formation, Pif1 was significantly enriched at tRNA genes, as previously described for a few tRNA loci (Figure S1A) (38). However, it was not enriched at sequences forming G4 or containing a G4 motif, contrary to what was reported in vegetatively growing cells (Figure S1A). We next compared the DSB-specific Pif1 signal (see Materials and Methods) with DSB hotspots, Mer3 peaks (36) and chromosome axis-associated Red1 peaks, which interact with DSB sites during recombination (39). As expected from our qPCR analyses, DSB-specific Pif1 signal correlated positively with DSB as well as with Mer3 signal (Figure 1B). Like Mer3, Pif1 peaks were mostly around DSB sites (Figure 1C and D). Pif1 was also weakly associated with axis-associated sites (Figure 1C and D), like many other DSB-related proteins (40). Conversely, Pif1 was not detected above background with DSB coldspots (Figure 1D, middle panel), suggesting that Pif1 is specifically recruited to DSB hotspots during meiosis. We wondered if Pif1 might be more required during DSB repair at sites containing tRNA genes or G4-prone sequences. However, we did not observe a stronger binding of Pif1 to DSB containing such structures (Figure S1B). Instead, Pif1 enrichment per DSB was rather uniform along the genome (Figure S1C). This indicates that Pif1 is likely part of the DSB repair machinery by homologous recombination, regardless of the presence of difficult to replicate DNA structures.

### Pif1 is required for longer gene conversions in *mlh2Δ*

Since Pif1 was associated with meiotic DSB sites, we tested whether it affected the length of recombination events. Due to the essential function of Pif1 in mitochondria, *pif1Δ* cells cannot sporulate. Therefore, we used a *pif1m2* allele, which has a point mutation at the second ATG of *PIF1* gene resulting in the loss of Pif1 from the nucleus while keeping a functional Pif1 in mitochondria (8). Indeed, in the SK1 background, *pif1m2* cells underwent meiosis normally and showed nearly wild-type spore viability (Figure S2A). To assess Pif1 function in recombination, we used S288C*SK1 *msh2Δ* hybrid diploids to sequence all the eight DNA strands generated after meiosis from one “octad” (Figure 2A). We sequenced meioses of *pif1m2* and *pif1m2 mlh2Δ* mutants and compared the results with our previous data that showed for the *mllh2Δ* mutant a significant increase in the length of CO- and NCO- associated heteroduplex DNA (hDNA) tracts (4).

**Figure 2:**
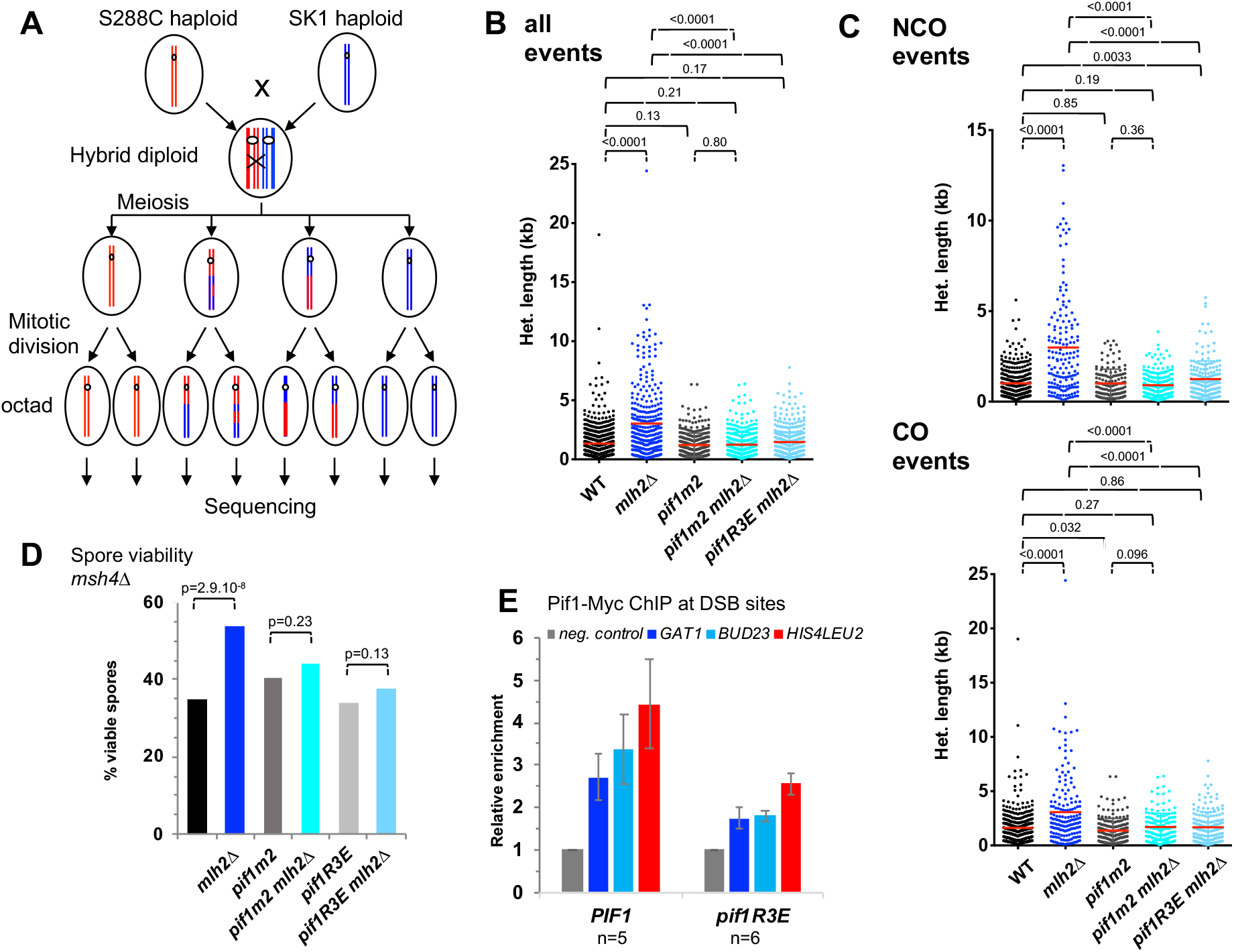
Pif1 is required for longer gene conversions in *mlh2Δ* through its interaction with PCNA. (A) Scheme of the experimental system to measure genome-wide recombination events. After meiosis, the four haploid spores of a tetrad are allowed to perform one mitosis and micromanipulated, in order to sequence DNA of the two daughters, allowing the recovery of the 8 DNA recombined strands (from one “octad”) from the initial diploid cell. (B) hDNA tracts lengths from meioses of the indicated genotype, all strains being also *msh2Δ*. The red horizontal bar indicates the mean value of pooled events from the meioses analyzed (4 meioses of WT and 2 meioses of each other relevant genotype). WT and *mlh2Δ* data are from (4). (C) Same as in (B) but for NCO-or CO-associated tracts. (B) and (C): Mann Whitney test p-values are indicated. (D) Effect of *mlh2Δ* and *pif1* mutants on spore viability of *msh4Δ* cells. All strains are *msh4Δ*. See also Table S2. Fisher’s exact test p-values are indicated. (E) Pif1-Myc levels at the three indicated meiotic DSB hotspots relative to a negative control site (LDLR) assessed by ChIP and qPCR at the 5 h time-point of meiotic time-courses. The signal is further normalized to the signal observed at t = 0 h. Values are the mean ± S.E.M. from the indicated number of independent experiments.

In the *mlh2Δ* mutant, the mean length of all hDNA tracts is 3.0 kb (2.2 median) as compared to 1.3 kb (1.0 median) in the reference (4). In *pif1m2*, hDNA tracts had a mean length of 1.2 kb (median 1.0), not different from the reference (Figure 2B). Strikingly, the *pif1m2 mlh2Δ* double mutant hDNA tracts had a mean length of 1.2 kb (median 1.0), similar to the reference and *pif1m2*, and significantly reduced compared to *mlh2Δ* (Figure 2B). The same behavior was observed when examining CO-associated and NCO-associated events separately, with one notable exception: for CO-associated events, *pif1m2* tracts had slightly reduced length (mean 1.4 kb, median 1.1 kb) compared to the reference (mean 1.6 kb, median 1.4kb), possibly revealing a minor function of Pif1 even in *MLH2* cells (Figure 2C). In addition, although the number of events was small, we noticed that whereas in the reference, there were 12 events longer than 5kb (4 meioses), there were only 2 in the *pif1m2* mutant (2 meioses) (Figure 2B). In this S288C*SK1 hybrid background, the single *mlh2Δ* and *pif1m2* mutants had reduced spore viability (Figure S2B). However, when combined together, the double mutant had no further reduction of spore viability, consistent with the suggestion that *pif1m2* mutation suppresses the defect of *mlh2Δ* (Figure S2B).

We previously showed that the longer hDNA tracts in the absence of MutLβ recruitment were accompanied by an increased spore viability of *zmm* mutants (4). Consistent with our octad results, we found that *pif1m2*, which had nearly no spore viability defect on its own, suppressed this increase in spore viability of *zmm* mutants, further confirming the specific function of Pif1 in the absence of MutLβ recruitment (Figure 2D). Next, we designed a separation-of-function mutant of *MLH2*, called *mlh2Δ500-536*, which specifically abolishes the interaction of Mlh2 with Mer3, but not with Mlh1 (Figure S3A-C). This mutant behaved exactly as *mlh2Δ* in terms of spore viability and suppression by Pif1 mutation (Figure S3D), indicating that the Mer3-MutLβ interaction is relevant for the effect of Pif1.

All these data show that Pif1 is not required to maintain a normal length of hDNA tracts, but is specifically involved in the formation of longer hDNA tracts in the absence of Mlh2. This is reminiscent to the function of Pif1 in BIR, and this prompted us to investigate if this function was linked to the stimulation of Pol δ.

### Pif1 interaction with PCNA plays an important role in longer gene conversions in *mlh2Δ*

Pif1 enhances DNA synthesis by Pol δ at least in part through its interaction with PCNA (16). We therefore investigated the *pif1R3E* mutant, which fails to interact with PCNA. The *pif1R3E* mutant cells sporulated and showed the same spore viability as wild type, confirming that other functions of Pif1 are maintained in this separation of function mutant (Figure S2A).

As expected from a role for the interaction with PCNA, the Pif1R3E mutant was less recruited to DSB sites than wild-type Pif1 (Figure 2E). Hence, we checked if the Pif1-PCNA interaction was important for longer recombination events in *mlh2Δ* cells. We analyzed patterns of hDNA of *pif1R3E mlh2Δ* double mutant in the S288C*SK1 *msh2Δ* hybrid. Like *pif1m2 mlh2Δ*, the *pif1R3E mlh2Δ* mutant showed hDNA tracts lengths (mean 1.5 kb, median 1.1) that were similar to the reference (mean 1.3 kb, median 1.0) (Figure 2B). When considering only the NCO events, *pif1R3E mlh2Δ* showed however slightly longer tracts than the reference (mean 1.2 versus 1.0 kb, median 1.1 versus 0.75 kb), meaning that the suppression of *mlh2Δ* may be partial (Figure 2C). However, in all cases, tracts were considerably shorter in the double *pif1R3E mlh2Δ* mutant compared to the single *mlh2Δ* (Figure 2B and C). The *pif1R3E* also decreased spore viability of *msh4Δ mlh2Δ*, as *pif1m2* (Figure 2D). Together, these data show that the direct interaction of Pif1 with PCNA is important for the generation of longer tracts observed in the absence of MutLβ.

### Mer3 interacts with Rfc1 and Pif1 *in vivo*

Mer3 belongs to the group of pro-crossover ZMM proteins, but our data show that it also acts earlier, as early as the D-loop intermediate stage, through its recruitment of MutLβ to meiotic DSB hotspots (present study and (4)). To identify its potential additional partners *in vivo*, we performed a TAP pulldown of Mer3 in synchronous meiotic cultures at 5 h in meiosis and analyzed pulled down proteins by quantitative label-free mass spectrometry, comparing Mer3-TAP and Mer3 untagged strains. In addition to the expected presence of Mlh1 and Mlh2 (MutLβ), we identified Rfc1, the subunit of RFC, the PCNA clamp loader, as a specific interactant (Figure 3A and Table S3). We confirmed this interaction by the reverse co-immunoprecipitation of Mer3-Flag by Rfc1-TAP (Figure 3B). This indicates that *in vivo*, Mer3 is present at the sites of DNA synthesis, where Rfc1 together with PCNA promotes D-loop extension by Pol δ (6). This suggests that Pif1 and Mer3 are present on recombination intermediates at the same stage. Although we did not detect Pif1 significantly enriched in our Mer3-TAP pull downs mass spectrometry analyses, we did detect Mer3 in Pif1-TAP pull downs by Western blot (Figure 3B), and confirmed this interaction by the reciprocal pull-down of Pif1 by Mer3-TAP (Figure 3C). These data are consistent with Mer3 acting during meiotic DNA repair synthesis, at the same sites and at the same stage as Pif1.

**Figure 3:**
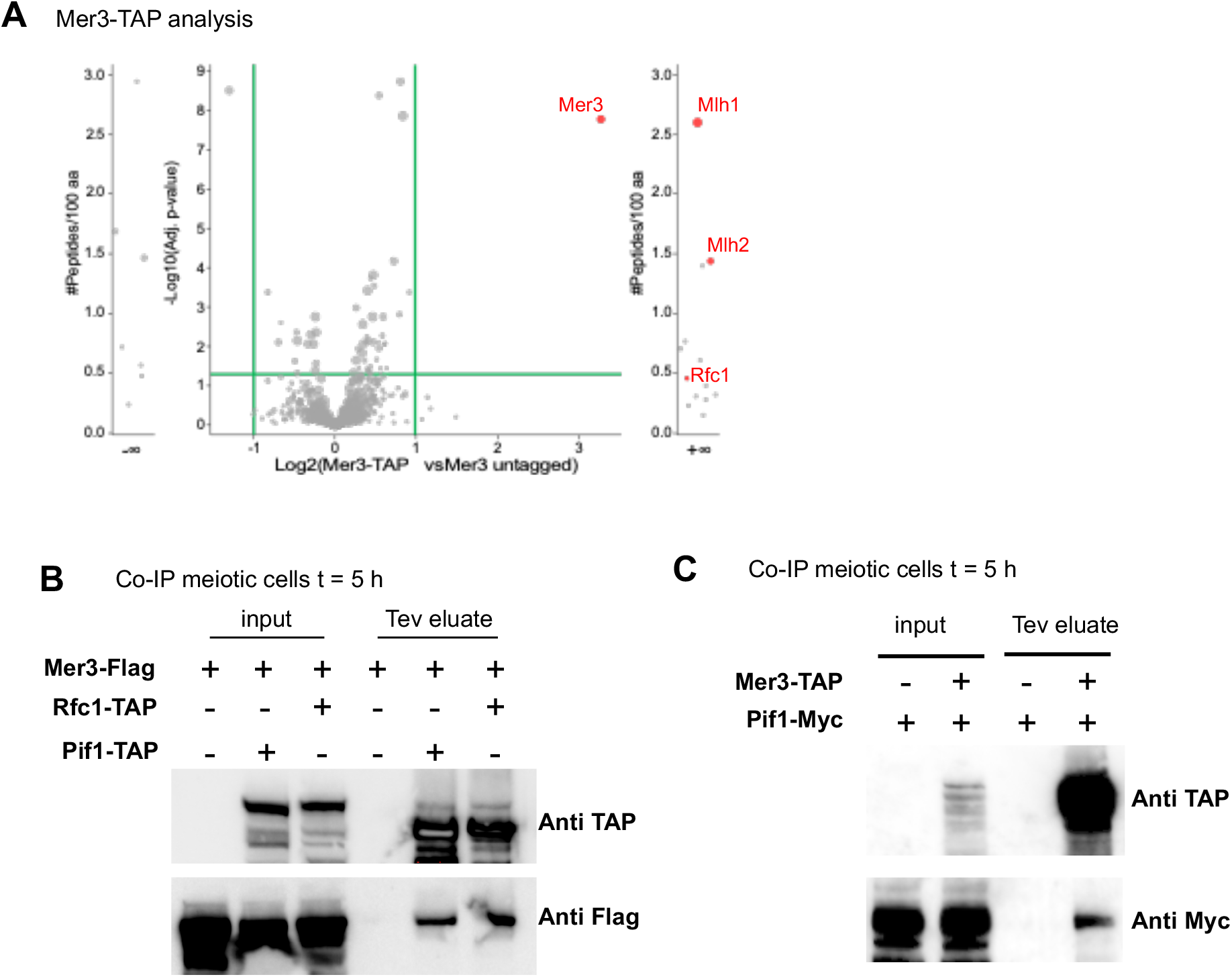
Mer3 interacts with the PCNA clamp loader Rfc1 and with Pif1 *in vivo*. (A) Volcano plot analysis identifying interactors of Mer3 in meiotic cells. Binding partners were obtained by using quantitative label-free mass spectrometry analysis of TAP pull down performed from four replicates. Shown are the fold changes (Mer3-TAP *versus* Mer3 untagged) quantified for proteins with 3 or more distinct peptides. Candidates are significantly enriched if fold change is higher than 2 and p-value is smaller than 0.05. External plots show proteins with peptides identified only in all replicates of one sample type, with 3 or more distinct peptides (left in Mer3 untagged, right in Mer3-TAP). See also Table S3. (B) Coimmunoprecipitation by Rfc1-TAP or Pif1-TAP from cells at 5 h in meiosis analyzed by Western blot. The Tev eluate produces a Rfc1-CBP or Pif1-CBP band smaller than the parental Rfc1-TAP and Pif1-TAP bands, respectively. (C) Coimmunoprecipitation by Mer3-TAP from cells at 5 h in meiosis analyzed by Western blot. The Tev eluate produces a Mer3-CBP band smaller than the parental Mer3-TAP band.

### The Mer3-MutLβ ensemble inhibits Pif1 *in vitro*

Pif1 is known to facilitate the formation of extension products by Pol δ from D-loop substrates (13). Using purified recombinant proteins and an assay similar to that used previously (Figure 4A and Figure S4A), we asked whether the Mer3-MutLβ ensemble inhibits Pif1 function *in vitro*. D-loops were formed with plasmid DNA and unlabeled ssDNA oligomer in the presence of purified Rad51, Rad54 and RPA (Figure S4B). D-loops were extended by Pol δ, RFC and PCNA, in the presence of labelled dCTP (Figure 4A). We then tested the effect of different combinations of Pif1, Mer3-hd (Mer3K167A helicase-dead mutant) and MutLβ on this reaction. Under the conditions used, in the absence of Pif1, only short extension products were seen (Figure 4A lane 2), while the addition of Pif1 resulted in longer extension products (Figure 4A lane 3). Alone, the addition of MutLβ at 3 or 30 nM concentration induced a modest decrease in D-loop extension products compared to Pif1 alone (Figure 4A lanes 4 and 5). A similar effect was seen upon the addition of Mer3-hd at 0.5 or 5 nM concentration (Figure 4A lanes 6 and 7). By contrast, when MutLβ and Mer3-hd were both used in the reaction, a significantly stronger reduction in the extension products was observed (Figure 4A lanes 8 and 9). These results suggest that the Mer3-MutLβ ensemble inhibits Pif1 *in vitro*.

**Figure 4:**
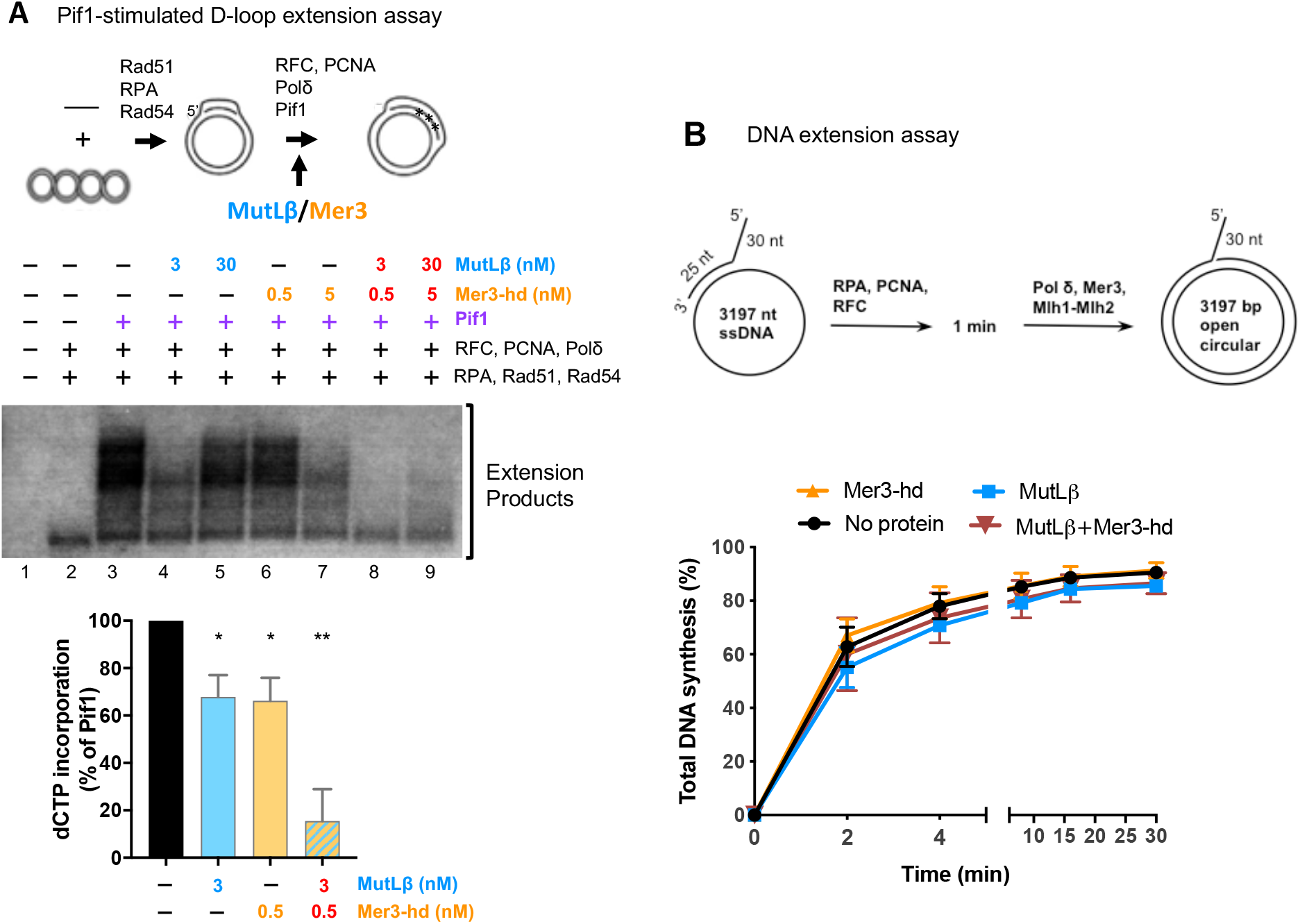
The Mer3-MutLβ ensemble inhibits Pif1 *in vitro*. (A) Pif1-stimulated D-loop extension by Pol δ. The experimental scheme for the D-loop extension assay from an unlabeled invading strand with [^32^P]-dCTP is adapted from (13). See also Figure S4. D-loop extension assay in the presence of the indicated combinations of Pif1, MutLβ (Mlh1-Mlh2) and Mer3-hd. Products were resolved by a native gel electrophoresis. A representative gel from four independent experiments is shown. The D-loop extension was quantified as percentage of the signal in the Pif1 lane of the relative experiment. The mean values ± S.E.M. are plotted from the four repeats. Statistical test: paired t test. **: p<0.01. *: p<0.05. (B) Kinetic analyses of primer extension by Pol δ in the presence of MutLβ or Mer3-hd. Quantification of the data are the mean values ± S.E.M. of three independent experiments. Representative gels are shown Figure S6.

To confirm that the inhibition of Pif1 by Mer3-MutLβ is specific and not related to mere binding of Mer3 and MutLβ to branched DNA substrates, we performed Electrophoretic Mobility Shift Assay (EMSA) with labelled D-loop structure in the presence of different combinations of Mer3 and MutLβ (Figure S5). As previously shown (4), Mer3 and MutLβ did bind to D-loop structures in a concentration-dependent manner. Importantly, at a concentration where a strong inhibition of Pif1 was seen in the D-loop extension assay (Figure 4A, line 8), Mer3 and MutLβ were not notably binding the DNA substrate alone in the EMSA assay (Figure S5, line 8) indicating the specific effect of Mer3-MutLβ on the D-loop extension actors. In addition, although Mer3 had a modest inhibitory effect on D-loop unwinding by Pif1, likely because of its affinity for this substrate, the addition of MutLβ did not further enhance this inhibition (Figure S6A). Finally, Mer3 and MutLβ, alone or in combination, had no effect on DNA extension by Pol δ (Figure 4B and S6B). Together, these experiments indicate that the Mer3-MutLβ ensemble specifically inhibits the Pif1-promoted DNA synthesis at D-loops.

## Discussion

We report here that Pif1 plays a role during DNA synthesis upon programmed meiotic break repair by homologous recombination. Our results thus reveal that Pif1 is not only required for BIR in vegetative cells, where one of the DSB ends is lost and a special replication bubble mediates DNA synthesis, but it is also involved during meiotic recombination events, especially when longer DNA synthesis by Pol δ takes place. This need for Pif1 is likely related to the fact that Pol δ, normally involved in lagging strand synthesis during S phase, does not interact with MCM, and therefore needs a helicase activity ahead to proceed over longer distances (6). We also found that Mer3-MutLβ restricts the Pol δ-mediated DNA synthesis by inhibiting Pif1, and therefore controls the length of meiotic heteroduplex DNA.

We found that Pif1 is recruited to programmed meiotic DSBs made by Spo11 (Figure 5, step (1)). Interestingly, Pif1 did not bind more at meiotic DSBs containing specific DNA features, such as G4 sequences or tRNA genes, proposed to be bound by Pif1 during DNA replication (37,38,41), suggesting that Pif1 is part of the normal machinery for DSB repair by homologous recombination. However, in otherwise wild-type cells, Pif1 only moderately contributes to the length of heteroduplex DNA. Its recruitment might therefore be a safeguard to allow DNA Pol δ to proceed, in case it encounters an obstacle such as DNA secondary structure or a R-loop, that would prevent the DNA synthesis required for DSB repair to take place. Our data reveal that specifically in meiotic cells, the activity of Pif1 at DSBs is nevertheless restrained, such that long DNA synthesis events, going over the size of resection tracts, are not permitted (Figure 5, left, step (2)).

**Figure 5:**
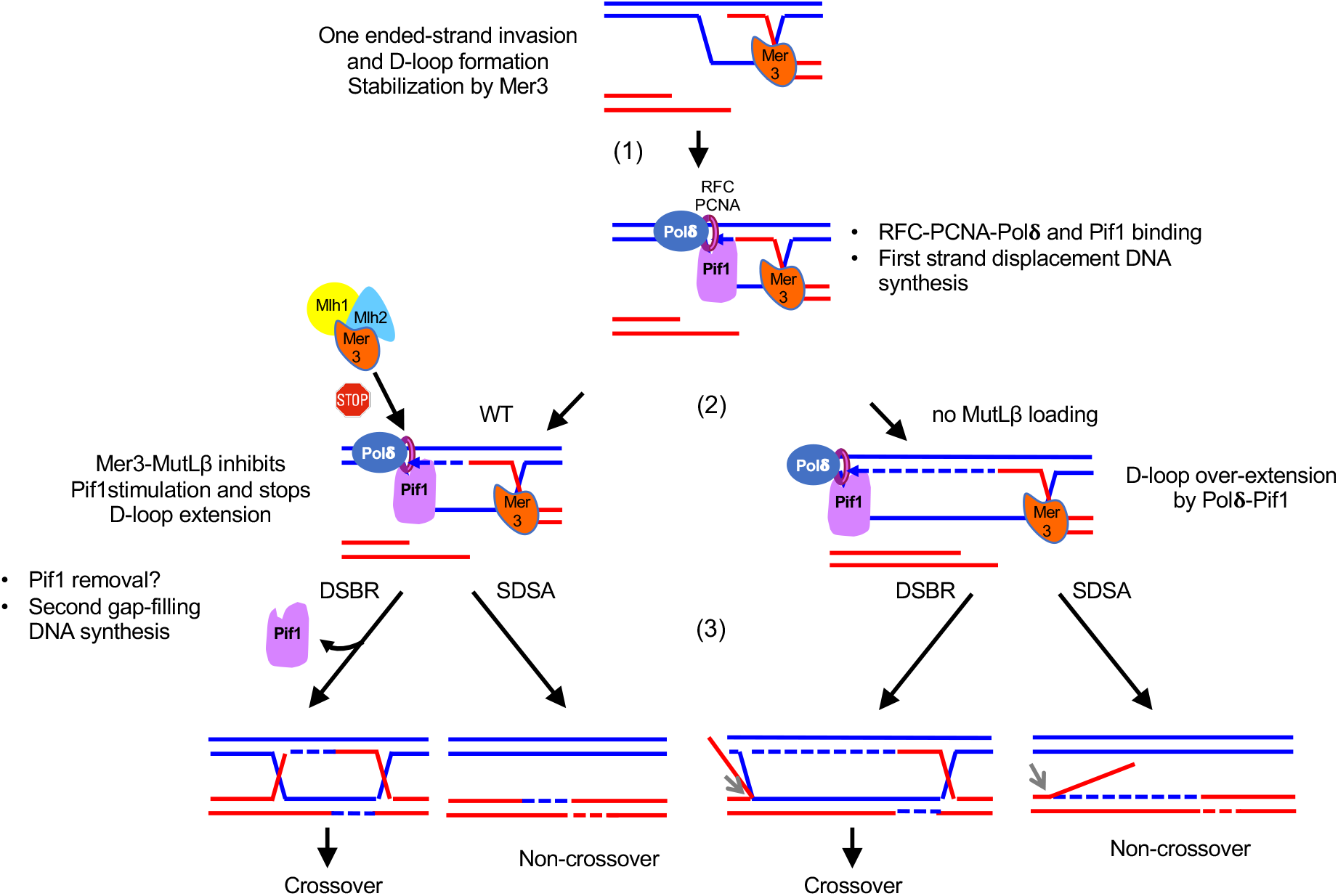
Model for the interplay between Pif1 and Mer3-MutLβ. Details are in the text.

During homologous recombination, DNA synthesis after end resection typically occurs in two steps, irrespective of a crossover or non-crossover outcome (Figure 5): at the first step, DNA synthesis occurs from the DSB end that invaded the homologous duplex and formed a D-loop intermediate. This step requires progressing through the D-loop and therefore involves strand-displacement (step (2)). At the second step, upon capture of the second end by the D-loop (DSBR) or first strand reannealing to the parental strand (SDSA), DNA synthesis operates through a gap filling mechanism (step (3)) (6). It was recently shown that for BIR, the two steps of DNA synthesis required employ Pol δ (14). In “normal” two ended-DSB repair by homologous recombination, although the first step is known to employ Pol δ, it is not known if the second does as well and if it could be influenced by the activity of Pif1 (6). From our genetic analyses of hDNA tracts, for the NCO events resulting from SDSA, we can unambiguously infer the tract of heteroduplex as resulting from first-end DNA synthesis (42), and these tracts are clearly much longer in the absence of Mer3-MutLβ, where they are promoted by Pif1 ((4) and this study). Unfortunately, the current way to analyze the tracts of DNA synthesis through the indirect measurement of hDNA tracts does not allow detection of second-end DNA synthesis during SDSA, nor distinction between first and second end synthesis for the CO events. However, we noticed that CO-associated hDNA tracts, which can result from first or second-end synthesis, are on average less increased than the NCO in the absence of Mer3-MutLβ (4). This would be compatible with the second end gap filling synthesis not being increased, and maybe not involving Pif1. Other experimental approaches, such as the direct detection of DNA synthesis from single events will be required to precisely determine the regulation of the two types of DNA synthesis. Finally, our biochemical experiments clearly indicate that Mer-MutLβ act specifically on Pif1-promoted DNA synthesis by Pol δ, which is also compatible with a specific effect on the first end DNA synthesis.

Based on our genetic and biochemical experiments, we therefore propose that in the absence of the Mer3-MutLβ control, Pif1 over-stimulates DNA synthesis during the first, strand-displacement, DNA synthesis (Figure 5, right, step (2)).

We can next envisage several modes of regulation by Mer3-MutLβ. First, the Mer3-MutLβ ensemble may be promoting second end capture or stand-reannealing. This would naturally stop the first end DNA synthesis by increasing the physical barrier to Pol δ progression. Therefore, in the absence of Mer3-MutLβ, DNA repair synthesis and second DSB end capture/re-annealing activity would be uncoupled and this would mimic a “BIR-like” situation, as if one DSB end was temporarily lost. However, this does not fit well with our *in vitro* data, which point to a more direct effect of Mer3-MutLβ on the strand-displacement activity of Pol δ stimulated by Pif1. As an alternative, since we Rfc1 is pulled-down with Mer3 from cells synchronously undergoing recombination, we propose that somehow, Mer3-MutLβ impede Pif1 action, possibly by competing for binding to RFC-PCNA. Binding of Mer3-MutLβ to RFC-PCNA would preclude Pif1 binding and stimulation. It would be interesting to determine how Mer3 interacts with RFC-PCNA, and why the presence of MutLβ is required with Mer3 to slow down DNA synthesis.

Besides its specific recruitment to meiotic DSBs, it is noteworthy that we did not detect any preferential binding of Pif1 to G4-prone sites in the genome in our meiotic samples. A possible explanation is that our highly synchronous meiotic cells are in a G2-like phase and do not experience DNA synthesis, contrary to cells in exponential phase where Pif1 occupancy was previously addressed (37). This is also consistent with models that propose that Pif1 unwinds G4-prone sequences during S phase, to prevent chromosome rearrangements (37,43). It would be interesting to test if removal of G4 by Pif1 occurs in the context of DNA synthesis, when Pif1 works with PCNA and Pol δ. For this, the *pif1R3E* mutant could be tested for its effect on the stability of a G4-containing minisatellite (22). At tRNA genes, we did detect a significant Pif1 enrichment in meiotic cells. It is possible, as suggested before, that at these sequences, Pif1 removes the R-loops formed because of replication/transcription collisions, outside or after S phase, prior to chromosome condensation and the first meiotic division (38).

Gene conversions resulting from the formation of heteroduplex are good to generate new allelic combinations, participating to genetic map distortions, but are also at risk of breaking favorable ancestral allele combinations (3). This may be why the generation of long gene conversion events is kept in check during meiosis, but it could also be to avoid the possibly mutagenic associated DNA synthesis.

## Data availability

Sequencing data were deposited at the NCBI Gene Expression Omnibus database with the accession numbers GSE164467 (Pif1-Myc13 ChIP-seq) and in the NCBI Sequence Read Archive under the accession number: SRP075437 (octad sequencing). Proteomic data were deposited on the Proteome Xchange via the PRIDE (Proteomics Identifications) partner repository with the identifiers PXD0023580 (Mer3-TAP analyses).

## Acknowledgments

We thank Judith Lopes for the *pif1m2* plasmid and Alberto Elias Villalobos for critical reading of the manuscript. We thank the Institut Curie NGS platform, supported by grants ANR-10-EQPX-03 and ANR10-INBS-09-08 and by the Cancéropôle Ile-de-France.

## Author Contributions

D.V. and V.B. conceived the project. D.V. and C.A. performed all yeast experiments. G.R. and L.R. performed all biochemical experiments under the supervision of P.C. F.D and D.L. performed the proteomic analysis. M.-C.M. and B.L. sequenced *pif3R3E* octads DNA and provided analysis tools for octad analyses. R.G. performed structure and interaction site predictions. V.B. supervised the project, and D.V. and V.B. wrote the manuscript with input from all the authors.

## Supplementary material

**Figure S1:**
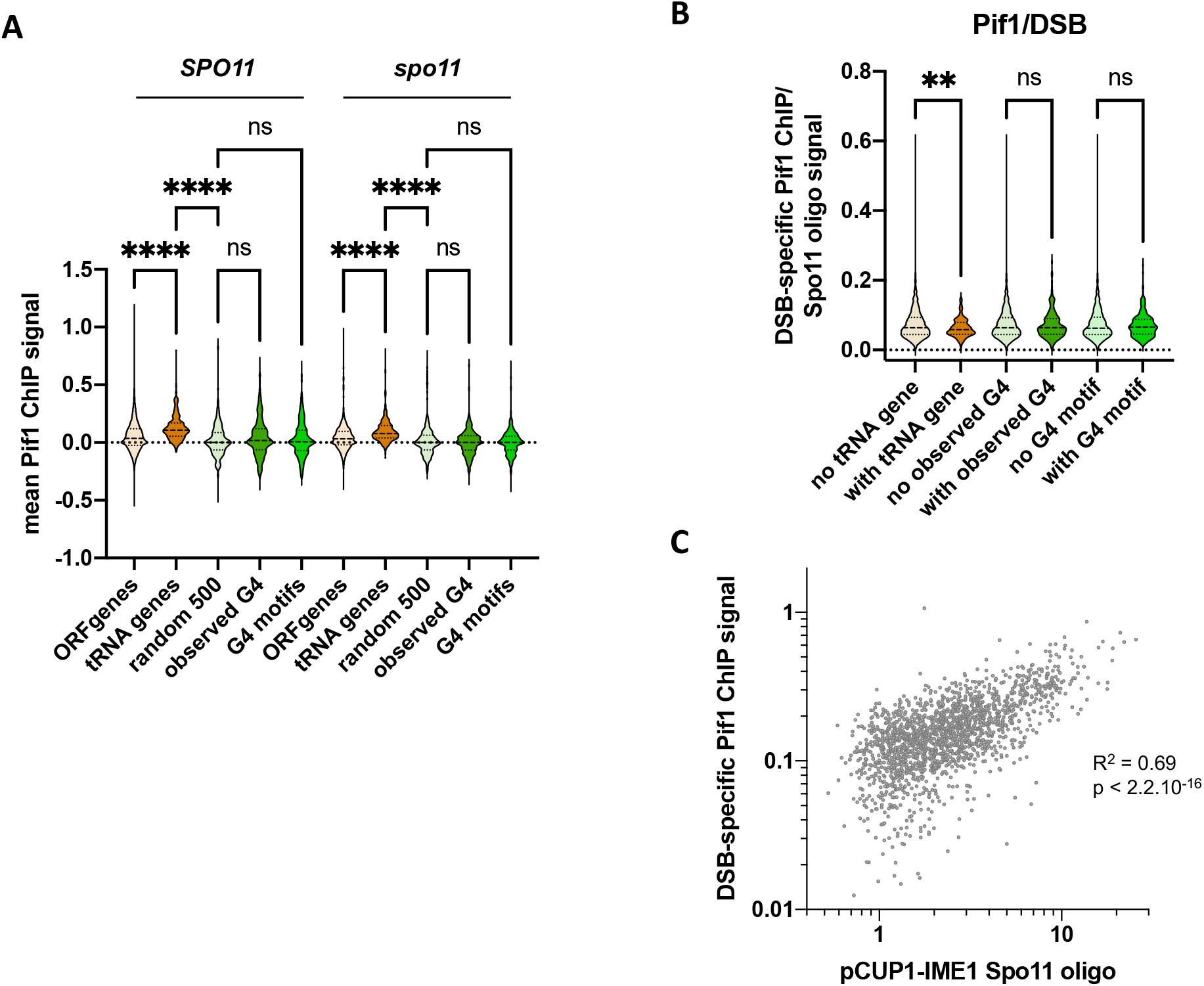
DSB-independent and DSB-specific Pif1 levels close to G4 sequences or tRNA genes. (A) The mean Pif1 ChIP-seq signal from both DSB-proficient (*SPO11*) and -deficient (*spo11*) meiotic cells was background-normalized and measured at 300 pb-long regions spanning the indicated genomic features. ORF genes corresponds to the yeast genome regions spanning from −100 bp to +200 bp from a TSS, to match the tRNA genes profile. Random 500 was generated by randomly selecting 500 genomic regions of 300 bp-long. Observed G4 are according to *in vivo* G4-seq signal measured in (1) (502 sites). G4 motifs are from (2) (668 sites). (B) The ratio of DSB-specific Pif1 ChIP-seq signal over *pCUP1-IME1* Spo11 oligo signal was computed on the width plus 1 kb on each side of the strongest 2000 Spo11 hotspots located in interstitial regions (more than 20 kb from a centromere and 40 kb from a telomere, 1829 hotspots). These were divided into hotspots containing or not the indicated feature: tRNA genes (96 DSB sites), observed G4 as defined in (1) (211 DSB sites), or G4 motifs as defined in (2) (271 DSB sites). (C) DSB-specific Pif1 ChIP-seq signal intensity as a function of DSB (p*CUP1-IME1* Spo11 oligo) signal intensity. The signal intensities at the locations comprised in the strongest 2000 Spo11 hotspots+1kb on each side were computed for each interstitial hotspot. The Pearson correlation coefficient and the associated p-value are indicated.

**Figure S2:**
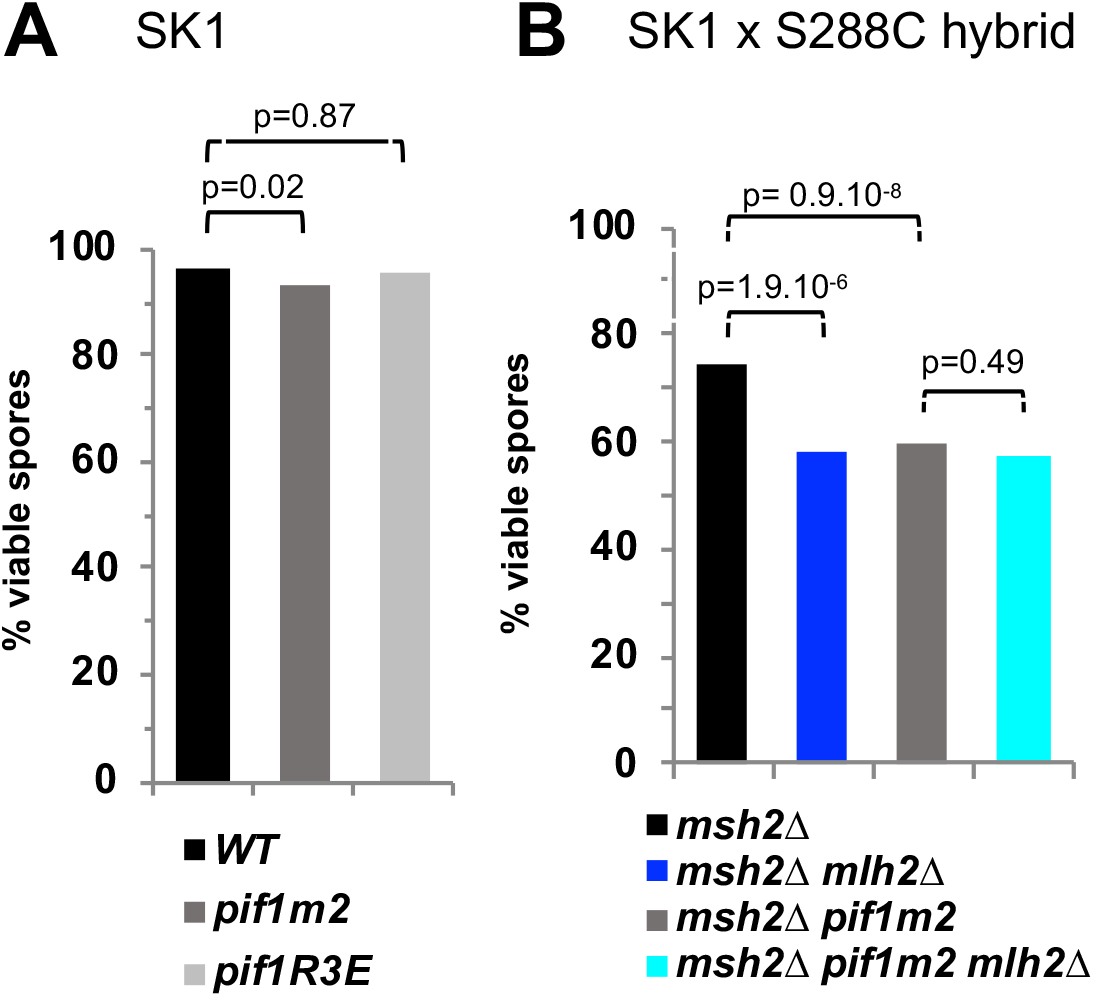
Effect of Pif1 mutants on spore viabilities in SK1 or hybrid strains. A) Spore viability of SK1 diploids with the indicated relevant genotype. (B) Spore viability of the SK1 × S288C hybrid used for octad analysis (See also Table S2)

**Figure S3:**
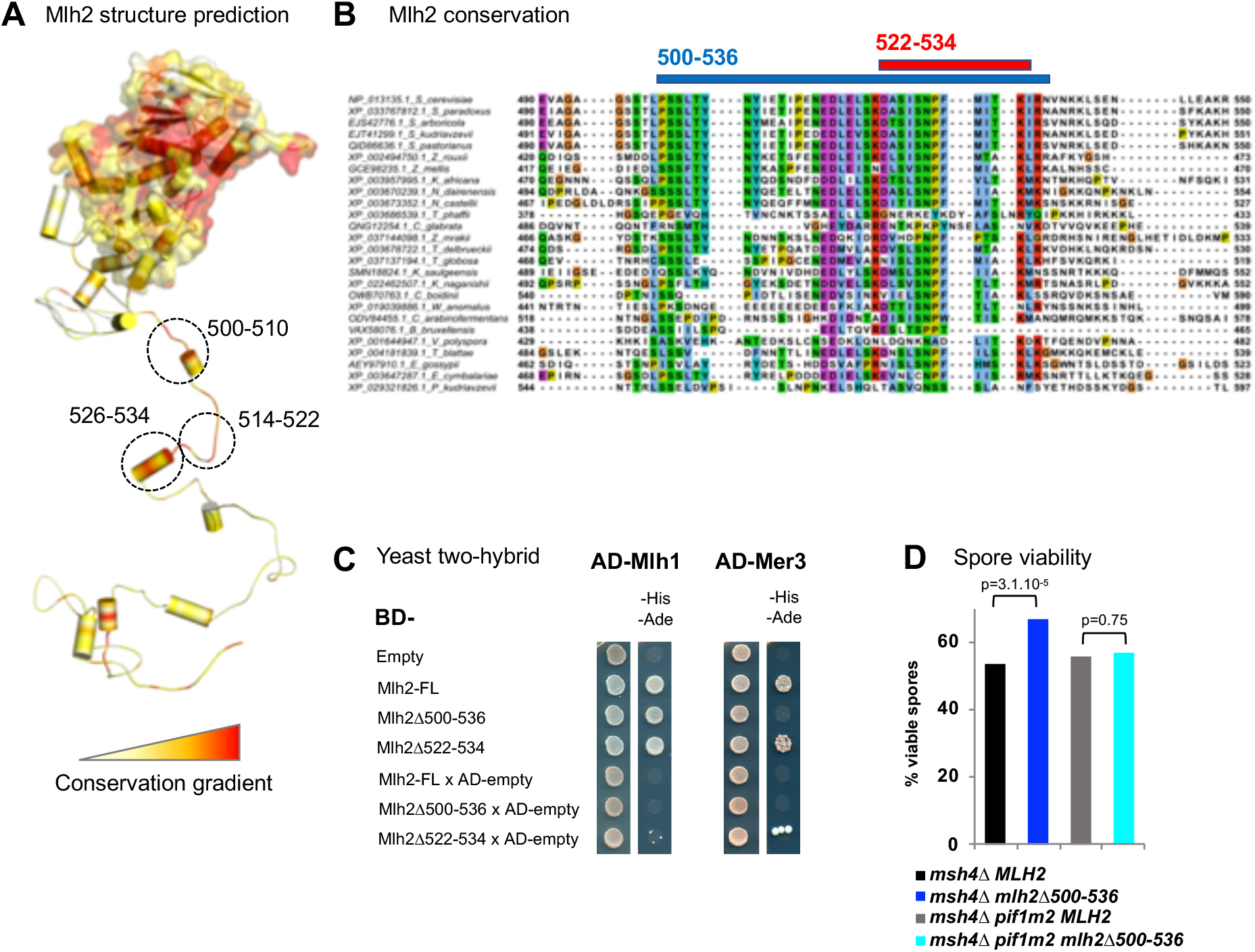
A separation of function mutant of *MLH2* deficient for interaction with Mer3 shows the same phenotype as an *mlh2Δ* mutant for *zmm* spore viability. (A) Model of Mlh2 structure. The candidate regions for interaction with Mer3 are indicated by circles. Conservation is indicated by a color code from red (conserved) to yellow (less conserved). (B) Alignment of several yeast species in the candidate Mer3 interacting region of Mlh2. (C) Two-hybrid interactions between Mlh2, Mlh1 and Mer3. The same number of cells of strains expressing the different fusion proteins (BD: Gal4 binding domain; AD: GAL4 activation domain) were plated on minimal media lacking the indicated aminoacids to select for interactions. Growth on -His-Ade medium indicates an interaction. The Mlh1-Mlh2 interaction is used as a positive control. FL: Full length. (D) Effect of *mlh2Δ500-536* and *pif1m2* mutants on spore viability of *msh4Δ* cells. See also Table S2. Fisher’s exact test p-values are indicated.

**Figure S4:**
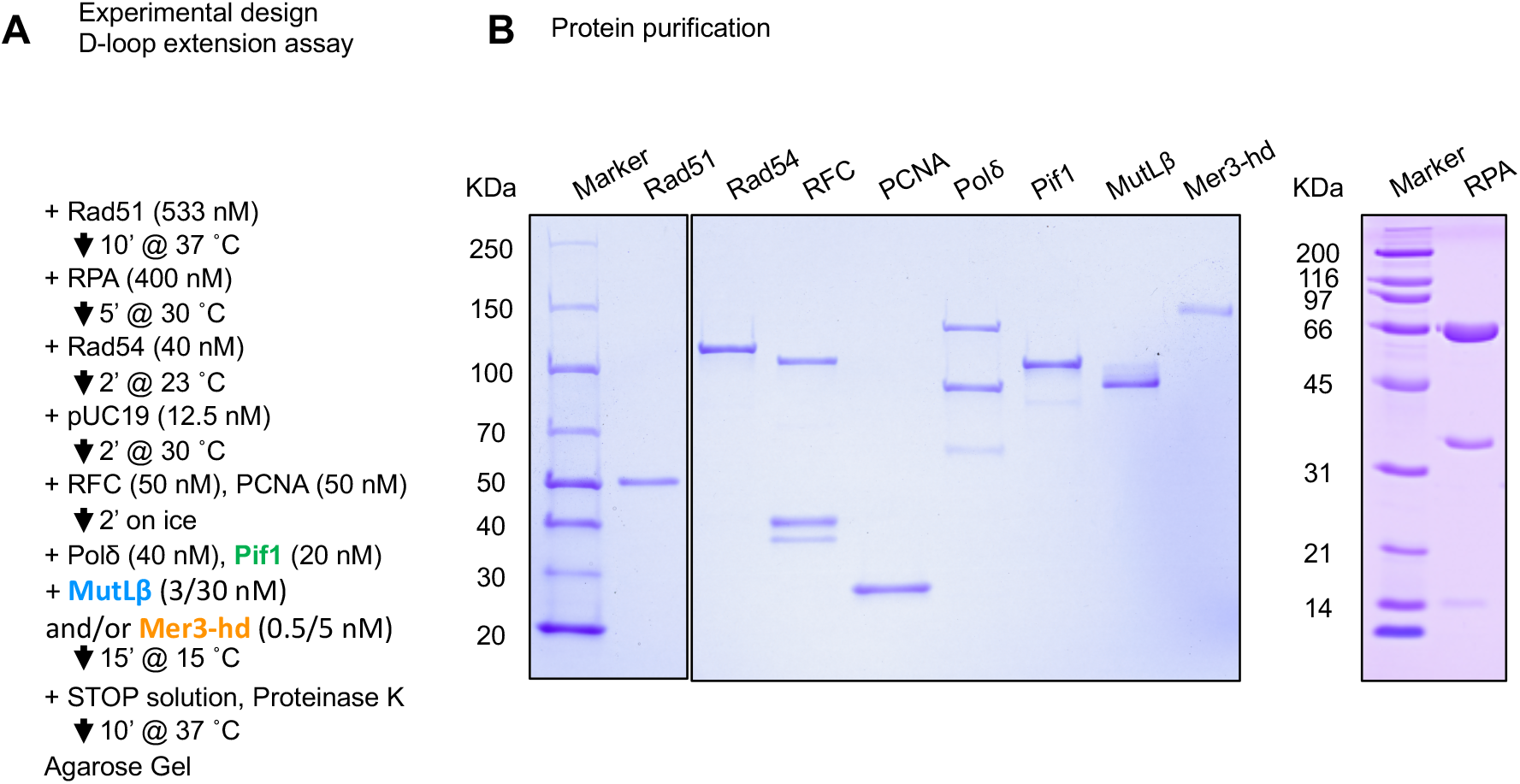
Experimental scheme for the D-loop extension assays and proteins used. (A) Experimental scheme for the D-loop extension assays. (B) Recombinant proteins were separated by SDS-PAGE on a 4-15% gradient gel and then visualized with Coomassie staining.

**Figure S5:**
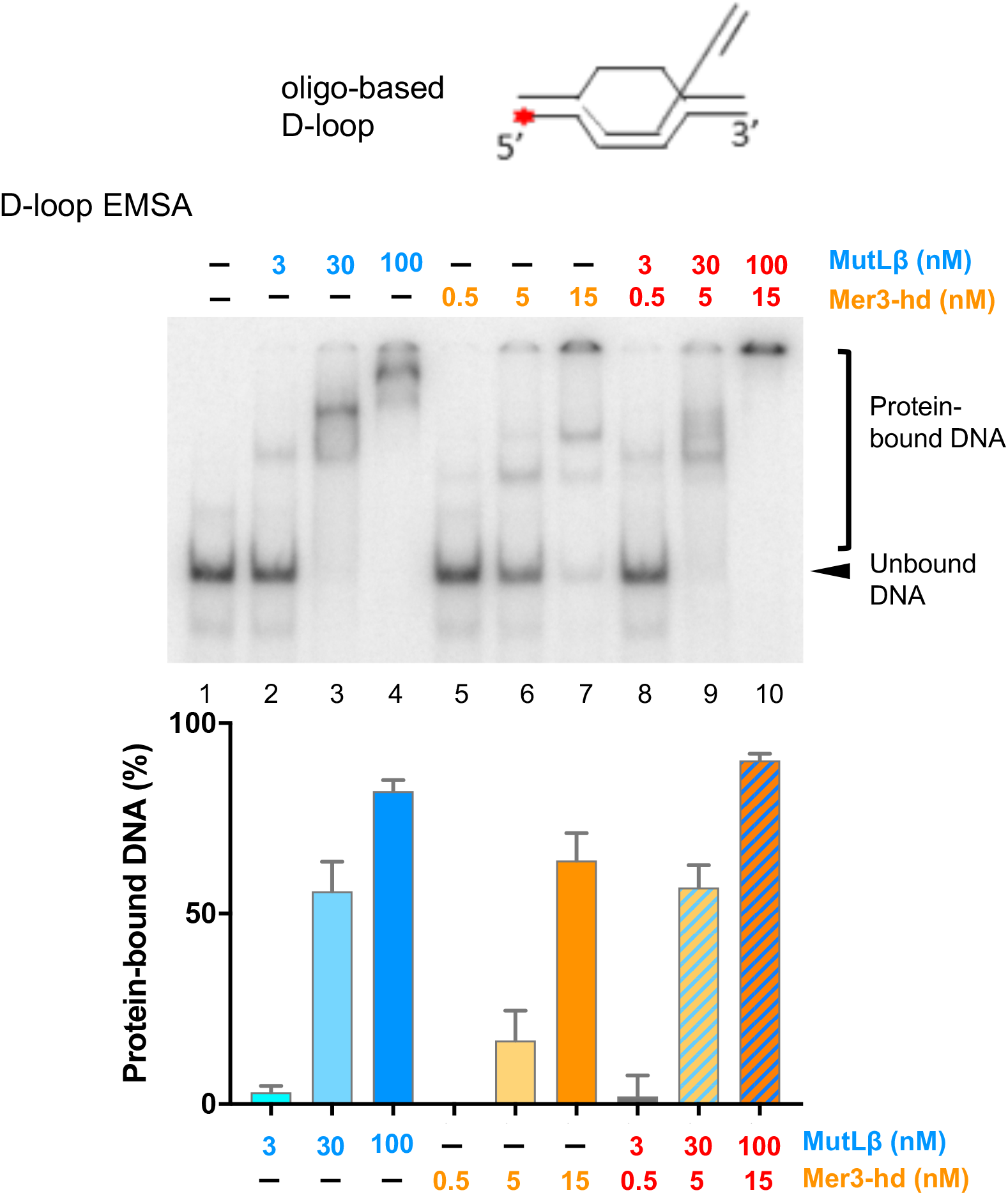
D-loop EMSA in the presence of the Mer3-hd and MutLβ. The cartoon illustrates the D-loop substrate used in the electrophoretic mobility shift assay (EMSA). Products were resolved by a native gel electrophoresis and quantified. A representative gel is shown. The mean values ± S.E.M. are plotted from three independent experiments.

**Figure S6:**
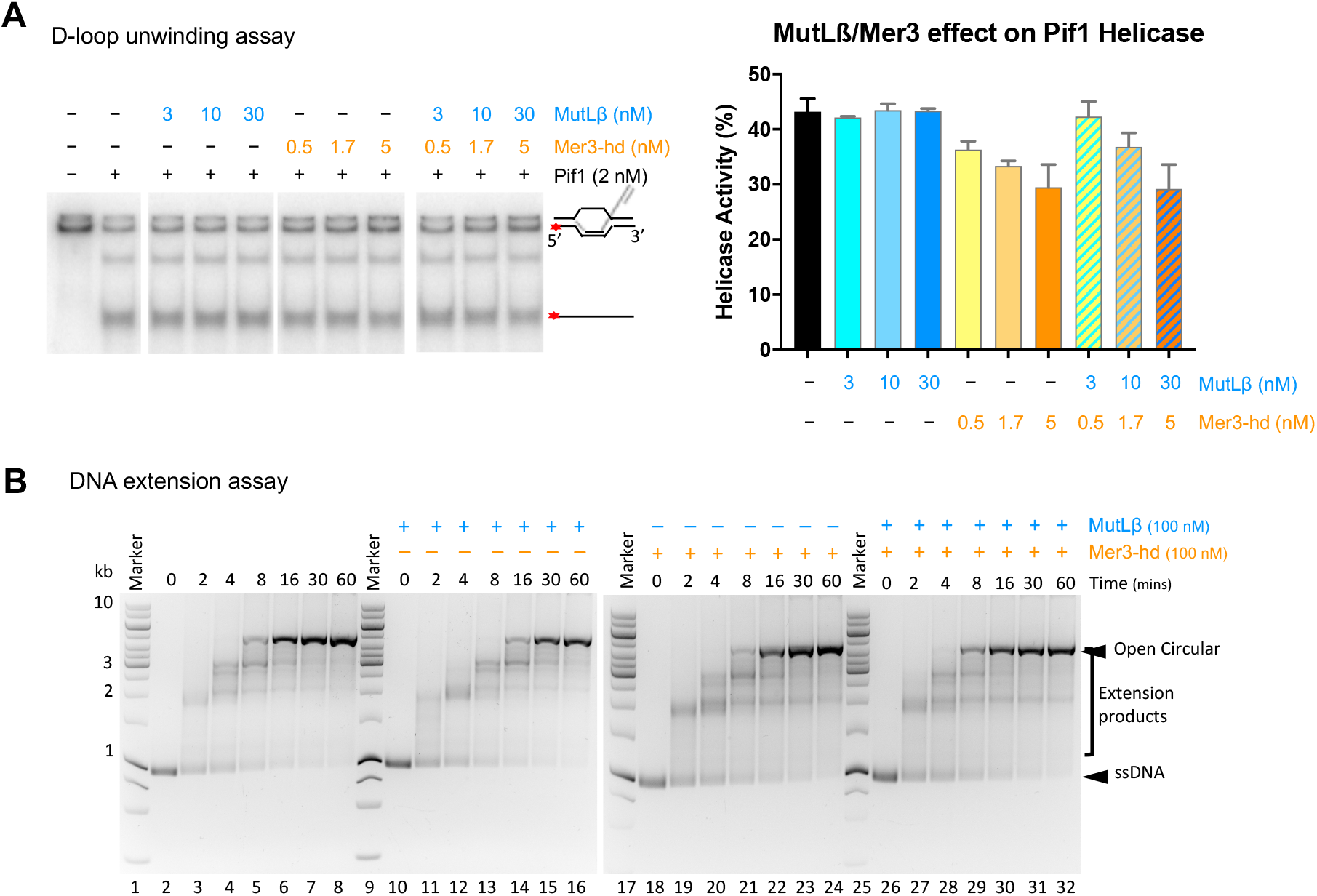
Effect of Mer3 and MutLβ on D-loop unwinding by Pif1 and on DNA extension by Pol δ. (A) D-loop unwinding assay. A representative experiment is shown, and the graph quantitation shows the mean ± S.E.M. of three independent experiments. (B) DNA extension by Pol δ. Representative **g**els from the experiments quantified in Figure 4B are shown.

**Table S1.**
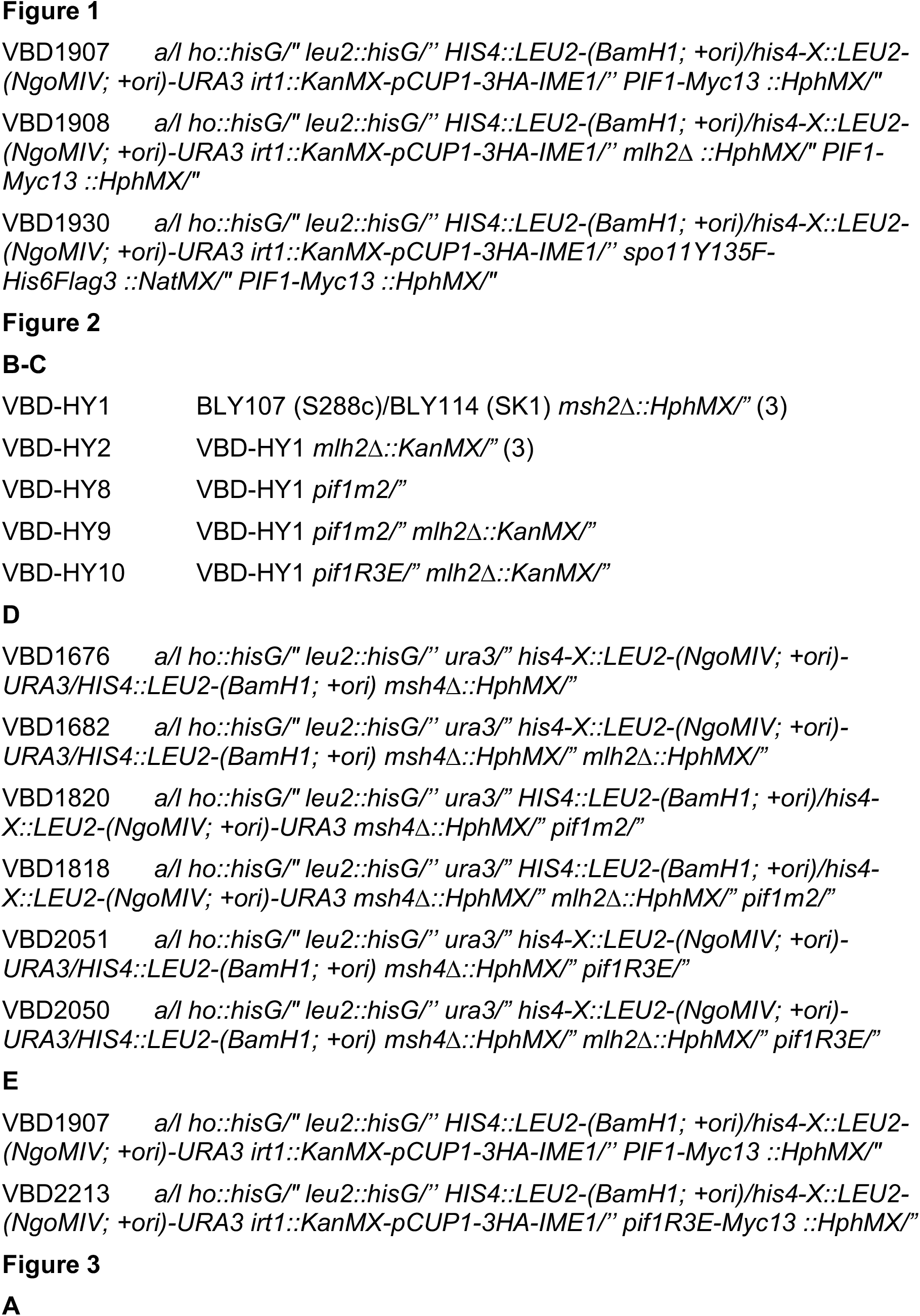
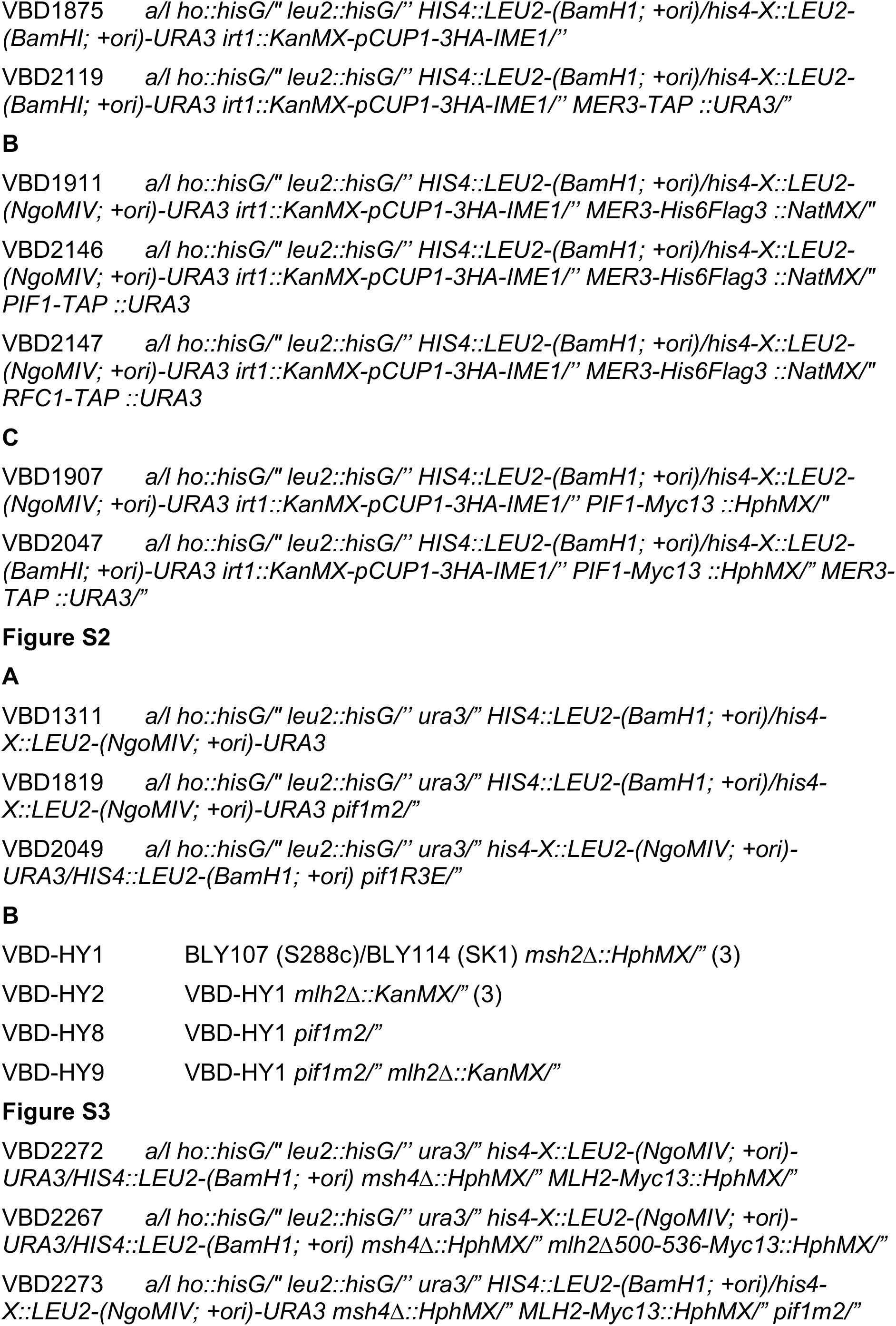
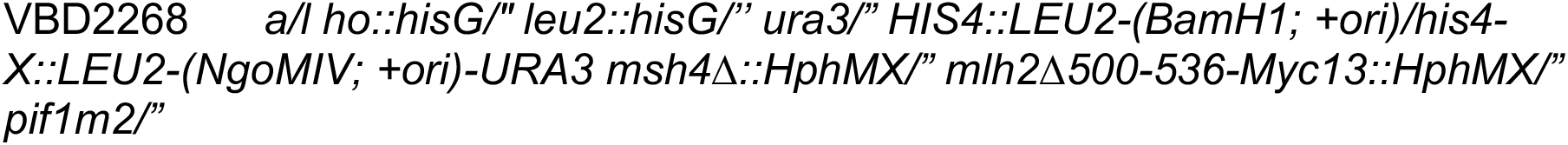
Strains used in this study. All strains are from the SK1 background, unless indicated otherwise.

**Table S2:** Spore viabilities.

| Strain name | Genotype | Tetrad type |  |  |  |  | Spore viability (%) | Number of tetrads | Fisher Test p-Value |
| --- | --- | --- | --- | --- | --- | --- | --- | --- | --- |
|  |  | 4 sp | 3 sp | 2 sp | 1 sp | 0 sp |  |  |  |
| VBD1311 | WT (3) | 131 | 9 | 7 | 0 | 0 | 96.1 | 147 | vs WT |
| VBD1819 | *pif1m2* | 104 | 21 | 6 | 0 | 1 | 93.0 | 132 | 0.0239 |
| VBD2049 | *pif1R3E* | 116 | 10 | 2 | 0 | 2 | 95.8 | 130 | 0.87 |
| VBD1676 | *msh4Δ* (3) | 27 | 3 | 24 | 3 | 63 | 35.0 | 120 | vs *msh4Δ* |
| VBD1682 | *msh4Δ mlh2Δ* (3) | 35 | 5 | 29 | 2 | 29 | 53.8 | 100 | 2.9E-08 |
| VBD1820 | *msh4Δ pif1m2* | 27 | 10 | 31 | 10 | 52 | 40.4 | 130 | vs *msh4Δ pif1m2* |
| VBD1818 | *msh4Δ pif1m2 mlh2Δ* | 36 | 6 | 29 | 10 | 49 | 44.2 | 130 | 0.23 |
| VBD2051 | *msh4Δ pif1R3E* | 40 | 6 | 45 | 13 | 103 | 33.9 | 207 | vs *msh4Δ pif1R3E* |
| VBD2050 | *msh4Δ pif1R3E mlh2Δ* | 50 | 7 | 41 | 9 | 101 | 37.5 | 208 | 0.13 |
| VBD2272 | *msh4Δ MLH2-Myc* | 42 | 4 | 20 | 3 | 35 | 53.6 | 104 | vs *msh4Δ MLH2-Myc* |
| VBD2267 | *msh4Δ mlh2Δ500-536-Myc* | 82 | 6 | 33 | 4 | 31 | 66.7 | 156 | 3.1E-05 |
| VBD2273 | *msh4Δ pif1m2 MLH2-Myc* | 33 | 17 | 21 | 6 | 27 | 55.5 | 104 | vs *msh4Δ pif1m2 MLH2-Myc* |
| VBD2268 | *msh4Δ pif1m2 mlh2Δ500-536-Myc* | 58 | 15 | 34 | 8 | 41 | 56.6 | 156 | 0.78 |
| HY1 | *msh2Δ* (3) | 48 | 33 | 26 | 13 | 0 | 74.2 | 120 | vs *msh2Δ* |
| HY2 | *msh2Δ mlh2Δ* (3) | 16 | 30 | 30 | 13 | 9 | 57.9 | 98 | 5.5E-07 |
| HY8 | *msh2Δ pifm2* | 32 | 19 | 40 | 16 | 11 | 59.5 | 118 | 1.9E-06 |
| HY9 | *msh2Δ mlh2Δ pif1m2* | 21 | 27 | 27 | 19 | 10 | 57.2 | 104 | 9.9E-08 |

## References

1. Hunter, N. (2015) Meiotic Recombination: The Essence of Heredity. Cold Spring Harb Perspect Biol, 7, a016618.

2. Pyatnitskaya, A., Borde, V. and De Muyt, A. (2019) Crossing and zipping: molecular duties of the ZMM proteins in meiosis. Chromosoma, 128, 181–198.

3. Cole, F., Keeney, S. and Jasin, M. (2012) Preaching about the converted: how meiotic gene conversion influences genomic diversity. Ann N Y Acad Sci, 1267, 95–102.

4. Duroc, Y., Kumar, R., Ranjha, L., Adam, C., Guerois, R., Md Muntaz, K., Marsolier-Kergoat, M.C., Dingli, F., Laureau, R., Loew, D. et al. (2017) Concerted action of the MutLbeta heterodimer and Mer3 helicase regulates the global extent of meiotic gene conversion. Elife, 6, e21900.

5. Kadyrov, F.A., Dzantiev, L., Constantin, N. and Modrich, P. (2006) Endonucleolytic function of MutLalpha in human mismatch repair. Cell, 126, 297–308.

6. McVey, M., Khodaverdian, V.Y., Meyer, D., Cerqueira, P.G. and Heyer, W.D. (2016) Eukaryotic DNA Polymerases in Homologous Recombination. Annu Rev Genet, 50, 393–421.

7. Wright, W.D., Shah, S.S. and Heyer, W.D. (2018) Homologous recombination and the repair of DNA double-strand breaks. J Biol Chem, 293, 10524–10535.

8. Schulz, V.P. and Zakian, V.A. (1994) The saccharomyces PIF1 DNA helicase inhibits telomere elongation and de novo telomere formation. Cell, 76, 145–155.

9. Boule, J.B., Vega, L.R. and Zakian, V.A. (2005) The yeast Pif1p helicase removes telomerase from telomeric DNA. Nature, 438, 57–61.

10. Chung, W.H. (2014) To peep into Pif1 helicase: multifaceted all the way from genome stability to repair-associated DNA synthesis. J Microbiol, 52, 89–98.

11. Donnianni, R.A. and Symington, L.S. (2013) Break-induced replication occurs by conservative DNA synthesis. Proc Natl Acad Sci U S A, 110, 13475–13480.

12. Saini, N., Ramakrishnan, S., Elango, R., Ayyar, S., Zhang, Y., Deem, A., Ira, G., Haber, J.E., Lobachev, K.S. and Malkova, A. (2013) Migrating bubble during break-induced replication drives conservative DNA synthesis. Nature, 502, 389–392.

13. Wilson, M.A., Kwon, Y., Xu, Y., Chung, W.H., Chi, P., Niu, H., Mayle, R., Chen, X., Malkova, A., Sung, P. et al. (2013) Pif1 helicase and Poldelta promote recombination-coupled DNA synthesis via bubble migration. Nature, 502, 393–396.

14. Donnianni, R.A., Zhou, Z.X., Lujan, S.A., Al-Zain, A., Garcia, V., Glancy, E., Burkholder, A.B., Kunkel, T.A. and Symington, L.S. (2019) DNA Polymerase Delta Synthesizes Both Strands during Break-Induced Replication. Mol Cell, 76, 371–381 e374.

15. Georgescu, R.E., Langston, L., Yao, N.Y., Yurieva, O., Zhang, D., Finkelstein, J., Agarwal, T. and O’Donnell, M.E. (2014) Mechanism of asymmetric polymerase assembly at the eukaryotic replication fork. Nat Struct Mol Biol, 21, 664–670.

16. Buzovetsky, O., Kwon, Y., Pham, N.T., Kim, C., Ira, G., Sung, P. and Xiong, Y. (2017) Role of the Pif1-PCNA Complex in Pol delta-Dependent Strand Displacement DNA Synthesis and Break-Induced Replication. Cell Rep, 21, 1707–1714.

17. Chia, M. and van Werven, F.J. (2016) Temporal Expression of a Master Regulator Drives Synchronous Sporulation in Budding Yeast. G3 (Bethesda), 6, 3553–3560.

18. Sanchez, A. and Borde, V. (2021) Methods to Map Meiotic Recombination Proteins in Saccharomyces cerevisiae. Methods in Molecular Biology, 2153, 295–306.

19. Puig, O., Caspary, F., Rigaut, G., Rutz, B., Bouveret, E., Bragado-Nilsson, E., Wilm, M. and Seraphin, B. (2001) The tandem affinity purification (TAP) method: a general procedure of protein complex purification. Methods, 24, 218–229.

20. Cannavo, E., Sanchez, A., Anand, R., Ranjha, L., Hugener, J., Adam, C., Acharya, A., Weyland, N., Aran-Guiu, X., Charbonnier, J.B. et al. (2020) Regulation of the MLH1-MLH3 endonuclease in meiosis. Nature.

21. Longtine, M.S., McKenzie, A.3rd,, Demarini, D.J., Shah, N.G., Wach, A., Brachat, A., Philippsen, P. and Pringle, J.R. (1998) Additional modules for versatile and economical PCR-based gene deletion and modification in Saccharomyces cerevisiae. Yeast, 14, 953–961.

22. Ribeyre, C., Lopes, J., Boule, J.B., Piazza, A., Guedin, A., Zakian, V.A., Mergny, J.L. and Nicolas, A. (2009) The yeast Pif1 helicase prevents genomic instability caused by G-quadruplex-forming CEB1 sequences in vivo. PLoS Genet, 5, e1000475.

23. Sanchez, A., Adam, C., Rauh, F., Duroc, Y., Ranjha, L., Lombard, B., Mu, X., Winterbert, M., Loew, D., Guarné, A. et al. (*in press*) Exo1 recruits Cdc5 polo kinase to MutLγ to ensure efficient meiotic crossover formation. Proc Natl Acad Sci U S A.

24. Serrentino, M.E., Chaplais, E., Sommermeyer, V. and Borde, V. (2013) Differential association of the conserved SUMO ligase Zip3 with meiotic double-strand break sites reveals regional variations in the outcome of meiotic recombination. PloS Genetics, 9, e1003416.

25. Sanchez, A., Adam, C., Rauh, F., Duroc, Y., Ranjha, L., Lombard, B., Mu, X., Wintrebert, M., Loew, D., Guarne, A. et al. (2020) Exo1 recruits Cdc5 polo kinase to MutLgamma to ensure efficient meiotic crossover formation. Proc Natl Acad Sci U S A, 117, 30577–30588.

26. De Muyt, A., Pyatnitskaya, A., Andreani, J., Ranjha, L., Ramus, C., Laureau, R., Fernandez-Vega, A., Holoch, D., Girard, E., Govin, J. et al. (2018) A meiotic XPF-ERCC1-like complex recognizes joint molecule recombination intermediates to promote crossover formation. Genes & development, 32, 283–296.

27. Zylicz, J.J., Bousard, A., Zumer, K., Dossin, F., Mohammad, E., da Rocha, S.T., Schwalb, B., Syx, L., Dingli, F., Loew, D. et al. (2019) The Implication of Early Chromatin Changes in × Chromosome Inactivation. Cell, 176, 182–197 e123.

28. Poullet, P., Carpentier, S. and Barillot, E. (2007) myProMS, a web server for management and validation of mass spectrometry-based proteomic data. Proteomics, 7, 2553–2556.

29. Valot, B., Langella, O., Nano, E. and Zivy, M. (2011) MassChroQ: a versatile tool for mass spectrometry quantification. Proteomics, 11, 3572–3577.

30. Anand, R., Pinto, C. and Cejka, P. (2018) Methods to Study DNA End Resection I: Recombinant Protein Purification. Methods Enzymol, 600, 25–66.

31. Biswas, E.E., Chen, P.H. and Biswas, S.B. (1995) Overexpression and rapid purification of biologically active yeast proliferating cell nuclear antigen. Protein Expr Purif, 6, 763–770.

32. Finkelstein, J., Antony, E., Hingorani, M.M. and O’Donnell, M. (2003) Overproduction and analysis of eukaryotic multiprotein complexes in Escherichia coli using a dual-vector strategy. Anal Biochem, 319, 78–87.

33. Johnson, R.E., Prakash, L. and Prakash, S. (2006) Yeast and human translesion DNA synthesis polymerases: expression, purification, and biochemical characterization. Methods Enzymol, 408, 390–407.

34. Levikova, M. and Cejka, P. (2015) The Saccharomyces cerevisiae Dna2 can function as a sole nuclease in the processing of Okazaki fragments in DNA replication. Nucleic Acids Res, 43, 7888–7897.

35. Opresko, P.L., Otterlei, M., Graakjaer, J., Bruheim, P., Dawut, L., Kolvraa, S., May, A., Seidman, M.M. and Bohr, V.A. (2004) The Werner syndrome helicase and exonuclease cooperate to resolve telomeric D loops in a manner regulated by TRF1 and TRF2. Mol Cell, 14, 763–774.

36. Sanchez, A., Adam, C., Rauh, F., Duroc, Y., Ranjha, L., Lombard, B., Mu, X., Wintrebert, M., Loew, D., Guarne, A. et al. (2020) Exo1 recruits Cdc5 polo kinase to MutLgamma to ensure efficient meiotic crossover formation. Proc Natl Acad Sci U S A.

37. Paeschke, K., Capra, J.A. and Zakian, V.A. (2011) DNA replication through G-quadruplex motifs is promoted by the Saccharomyces cerevisiae Pif1 DNA helicase. Cell, 145, 678–691.

38. Tran, P.L.T., Pohl, T.J., Chen, C.F., Chan, A., Pott, S. and Zakian, V.A. (2017) PIF1 family DNA helicases suppress R-loop mediated genome instability at tRNA genes. Nat Commun, 8, 15025.

39. Sun, X., Huang, L., Markowitz, T.E., Blitzblau, H.G., Chen, D., Klein, F. and Hochwagen, A. (2015) Transcription dynamically patterns the meiotic chromosome-axis interface. Elife, 4.

40. Panizza, S., Mendoza, M.A., Berlinger, M., Huang, L., Nicolas, A., Shirahige, K. and Klein, F. (2011) Spo11-accessory proteins link double-strand break sites to the chromosome axis in early meiotic recombination. Cell, 146, 372–383.

41. Rodriguez, R., Miller, K.M., Forment, J.V., Bradshaw, C.R., Nikan, M., Britton, S., Oelschlaegel, T., Xhemalce, B., Balasubramanian, S. and Jackson, S.P. (2012) Small-molecule-induced DNA damage identifies alternative DNA structures in human genes. Nat Chem Biol, 8, 301–310.

42. Marsolier-Kergoat, M.C., Khan, M.M., Schott, J., Zhu, X. and Llorente, B. (2018) Mechanistic View and Genetic Control of DNA Recombination during Meiosis. Molecular Cell, 70, 9–20 e26.

43. Lopes, J., Piazza, A., Bermejo, R., Kriegsman, B., Colosio, A., Teulade-Fichou, M.P., Foiani, M. and Nicolas, A. (2011) G-quadruplex-induced instability during leading-strand replication. EMBO J, 30, 4033–4046.

44. Zhu, X. and Keeney, S. (2015) High-Resolution Global Analysis of the Influences of Bas1 and Ino4 Transcription Factors on Meiotic DNA Break Distributions in Saccharomyces cerevisiae. Genetics, 201, 525–542.

## Supplemental references

1. Marsico, G., Chambers, V.S., Sahakyan, A.B., McCauley, P., Boutell, J.M., Antonio, M.D. and Balasubramanian, S. (2019) Whole genome experimental maps of DNA G-quadruplexes in multiple species. Nucleic Acids Res, 47, 3862–3874.

2. Capra, J.A., Paeschke, K., Singh, M. and Zakian, V.A. (2010) G-quadruplex DNA sequences are evolutionarily conserved and associated with distinct genomic features in Saccharomyces cerevisiae. PLoS Comput Biol, 6, e1000861.

3. Duroc, Y., Kumar, R., Ranjha, L., Adam, C., Guerois, R., Md Muntaz, K., Marsolier-Kergoat, M.C., Dingli, F., Laureau, R., Loew, D. et al. (2017) Concerted action of the MutLbeta heterodimer and Mer3 helicase regulates the global extent of meiotic gene conversion. Elife, 6, e21900.

